# Mitoxantrone Targets Both Host and Bacteria to Overcome Vancomycin Resistance in *Enterococcus faecalis*

**DOI:** 10.1101/2022.10.24.513631

**Authors:** Ronni A. G. da Silva, Jun Jie Wong, Haris Antypas, Pei Yi Choo, Karlyn Goh, Shreya Jolly, Cui Liang, Leona Tay Kwan Sing, Mark Veleba, Guangan Hu, Jianzhu Chen, Kimberly A. Kline

## Abstract

Among Enterococci, intrinsic and acquired resistance to antibiotics such as β-lactams and vancomycin critically limit treatment options for infection with these opportunistic pathogens. Antimicrobials that enhance the host immune response are emerging as alternative approaches, with the potential to overcome bacterial resistance. Here, we investigate the antibiotic and immunological activity of the anticancer agent mitoxantrone (MTX) *in vitro* and *in vivo* against vancomycin resistant *Enterococcus faecalis* (VRE). We show that, *in vitro*, MTX is a potent antibiotic against Gram-positive bacteria with a minimal inhibitory concentration (MIC) of ~1 μg/ml through induction of reactive oxygen species and DNA damage. MTX synergises with vancomycin and lowers the vancomycin concentration required to kill VRE by over 140-fold. This synergy is specific to vancomycin-resistant, but not susceptible strains because vancomycin rendered the resistant strains more permeable to MTX and thus MTX-mediated DNA damage. In a murine wound infection model, MTX treatment effectively reduced VRE bacterial numbers by 120-fold and with further reduction when combined with vancomycin. Wounds treated with MTX had significantly higher numbers of macrophages and higher pro-inflammatory cytokines compared to untreated wounds. In addition, MTX augmented intracellular bacterial killing by both murine and human macrophages by upregulating the expression of lysosomal hydrolases cathepsins D and H, and β-Hexosaminidase. These results show that MTX is a potent antibiotic against Gram-positive bacteria, synergizes with vancomycin, enhances macrophage recruitment and intracellular bactericidal activity, and represents a promising dual bacterium- and host-targeted therapeutic for overcoming vancomycin resistance.

**One sentence summary:** Mitoxantrone synergizes with vancomycin against vancomycin resistant bacterial strains via direct antibiotic activity and by augmenting both host macrophage recruitment to the site of infection and macrophage bactericidal activity.

## Introduction

Antibiotic resistance represents a major global health threat. Recent estimates attribute 4.95 million deaths in 2019 to antimicrobial resistance (AMR) (*1*), with this number predicted to climb to 10 million deaths annually by 2050 (*2*). Thus, multi-pronged treatment approaches, including antimicrobials that overcome existing resistance mechanisms and host-directed adjuvant therapies that enhance natural immune responses are emerging as important alternatives to fight bacterial infections (*3*).

*Enterococcus faecalis, Staphylococcus aureus*, and *Pseudomonas aeruginosa are* among the most frequently isolated bacterial species from wounds including burns, diabetic foot ulcers, surgical sites, and chronic wounds (*4–7*). *E. faecalis* possess intrinsic resistance to penicillin, ampicillin, and cephalosporins (*8*). An estimated ~30% of healthcare associated enterococcal infections are caused by vancomycin-resistant strains, further reducing treatment options (*9*). Vancomycin-resistant enterococci (VRE) are on the Centers for Disease Control and Prevention (CDC) serious threat watch list, with $539 million in estimated attributable healthcare costs in 2017 alone (*9*). This public health threat will continue to grow as enterococci acquire resistance to last resort antimicrobials such as quinupristin-dalfopristin, linezolid, daptomycin, and tigecycline (*8, 10*).

Understanding the molecular mechanism behind resistance is fundamental to develop new strategies to fight it. Vancomycin inhibits Gram-positive bacteria by targeting cell wall synthesis (*11*). In vancomycin resistant strains, proteins of a two-component regulatory system (such as VanRS) sense the binding of vancomycin to peptidoglycan (PG) precursor D-alanyl-D-alanine (D-Ala-D-Ala) dipeptide termini. Cell wall disturbances trigger the PG precursor replacement to D-alanyl-D-lactate (D-Ala-D-Lac) by other proteins encoded by the *van* operon, impeding vancomycin binding. Importantly, the D,D-carboxypeptidase VanY removes the terminal D-Ala residue from PG in the cell wall, and the enzyme VanX hydrolyzes D-Ala-D-Ala, thereby reducing the pool of available D-Ala-D-Ala (*10, 12*) that vancomycin could bind to.

In addition to intrinsic and acquired AMR that complicate the treatment of enterococcal infection, we have previously shown that extracellular *E. faecalis* can subvert immune activation during infection (*13*). Moreover, *E. faecalis* can be internalized by keratinocytes and macrophages, alter endo-lysosomal trafficking, and replicate intracellularly leading to a hyper-infective phenotype (*14*). Furthermore, persistent *E. faecalis* wound infection is associated with lowered cytokine levels in the wound which could impair wound healing (*15*). Thus, infections with *E. faecalis* are perfectly positioned to benefit from host-targeted immunotherapies that may counteract the immune-modulating action of this microbe.

One potential target for host-directed immunotherapy is to promote enterococcal clearance by macrophages. Following phagocytosis, macrophages generate reactive oxygen and nitrogen species (*16, 17*), mobilize transition metals to intoxicate microbial cells (*18*), and acidify the phagosome to activate lysosomal enzymes to eliminate internalized bacteria (*19*). Macrophages also present pathogen antigens to other immune cells and secrete cytokines to recruit immune cells to the infected site (*20*). Moreover, macrophages possess high transcriptional plasticity which allow them to polarize and change their phenotypic profile depending on host or external stimuli (*21, 22*). Macrophage polarization can exist on a spectrum, including macrophages associated with a pro-inflammatory state in response to infection, and macrophages linked to tissue repair and remodelling (*23*). Altogether, due to their plasticity and pivotal role in fighting infections and promoting tissue repair, drugs that can reprogram macrophages toward an optimal response may improve infection outcomes.

Repurposed compounds offer an excellent opportunity in the pursuit of new therapies that can target both the host and the pathogen. The vast knowledge and safety validation of drugs deployed to treat other health conditions can drastically reduce the time and cost in the development of new therapeutical approaches (*24*). From a pool of 4126 compounds (including 760 FDA-approved drugs), we previously identified a series of compounds as capable of re-programming macrophages into a pro-inflammatory state or anti-inflammatory state (*25*). In this study, we evaluated selected compounds for antibiotic activity and macrophage augmentation leading to enhanced bacterial clearance. Here we report that mitoxantrone (MTX), an antineoplastic agent commonly used to treat acute leukaemia, prostate, and breast cancer, as well as multiple sclerosis by disrupting DNA replication in mammalian cells, is both antimicrobial and immunomodulatory. Our data show that MTX both resensitizes VRE to killing by vancomycin and promotes macrophage recruitment and activation, rendering MTX an attractive candidate for difficult to treat VRE infections.

## Results

### MTX exhibits potent antibiotic activities *in vitro* and *in vivo*

We evaluated 18 compounds that were previously shown to re-program macrophages (*26*) for their ability to enhance macrophage-mediated killing of intracellular bacteria *in vitro*. MTX was among the most potent compounds **(fig. S1, table S1)** and we therefore tested its antibiotic activity by measuring minimal inhibitory concentration (MIC) against a panel of both Gram-positive and Gram-negative bacterial strains in the absence of macrophages. Bacteria were grown in increasing concentrations of MTX, ranging from 0.4 μg/ml to 51.2 μg/ml, for 24 hours (h). MTX was 10-20-fold more potent in inhibiting the growth of Gram-positive bacteria (~1 μg/ml MIC) compared to Gram-negative bacteria **(Table 1)**. Among the Gram-positive bacteria, MTX similarly inhibited the growth of vancomycin-resistant *E. faecalis* V583 (our prototypic VRE strain going forward), *E. faecium* AUS0004 and *E. faecium* E745, and vancomycin-sensitive *E. faecalis* OG1RF and *S. aureus* USA300 (MRSA).

**Table 1.**
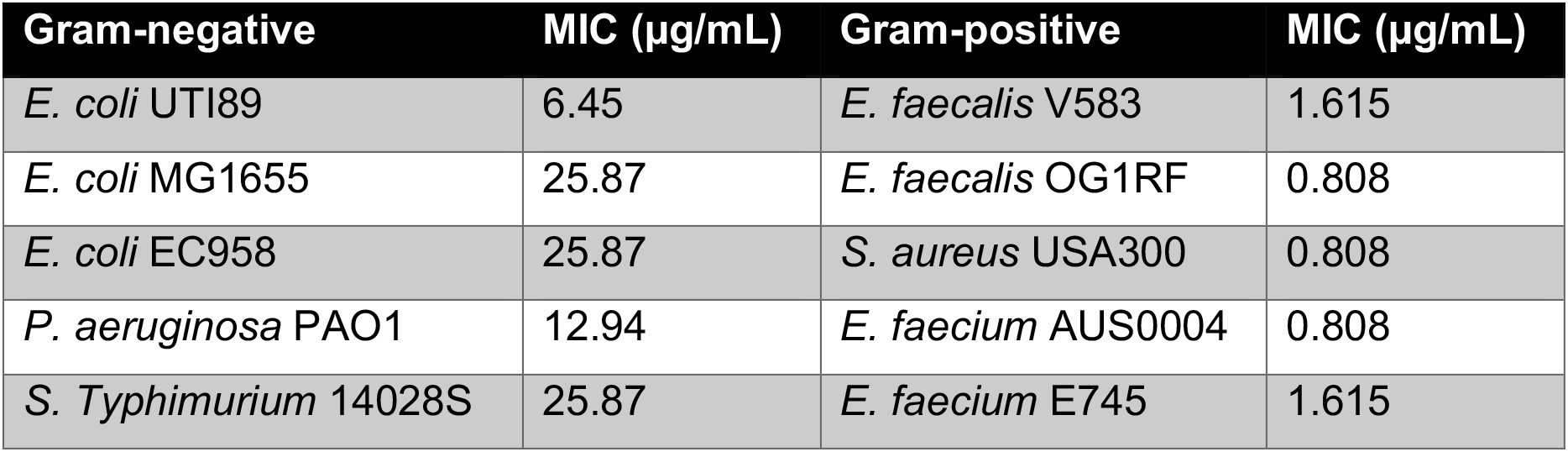
MTX is more efficient against Gram-positive bacteria in DMEM. MTX MIC for different bacterial species.

We tested the potency of MTX in inhibiting bacterial growth *in vivo* using a mouse wound infection model (*15*). Wounds were infected with 10^7^ colony forming units (CFU) of VRE, followed with addition of 10 μl of MTX (0.515 μg/mL) or PBS. Twenty-four hours post infection (hpi), the median CFU per wound was 6 x 10^8^ for PBS controls, whereas the median CFU was 4.9 x 10^6^ for MTX-treated wounds, a reduction of ~120-fold (**Fig. 1A**). Similarly, MTX treatment reduced the median CFU of MRSA USA300 and *P. aeruginosa* PAO1 by ~60 and ~3.5 fold **(Fig. 1B-C)**, respectively. Moreover, infected wounds presented with a purulent exudate which was visibly reduced with MTX treatment, most prominently for VRE and *S. aureus* infection **(Fig. 1D).** These results show that MTX exhibits potent antibiotic activity both *in vitro* and *in vivo*, especially against Gram-positive bacterial species, including those resistant to vancomycin.

**Fig. 1.**
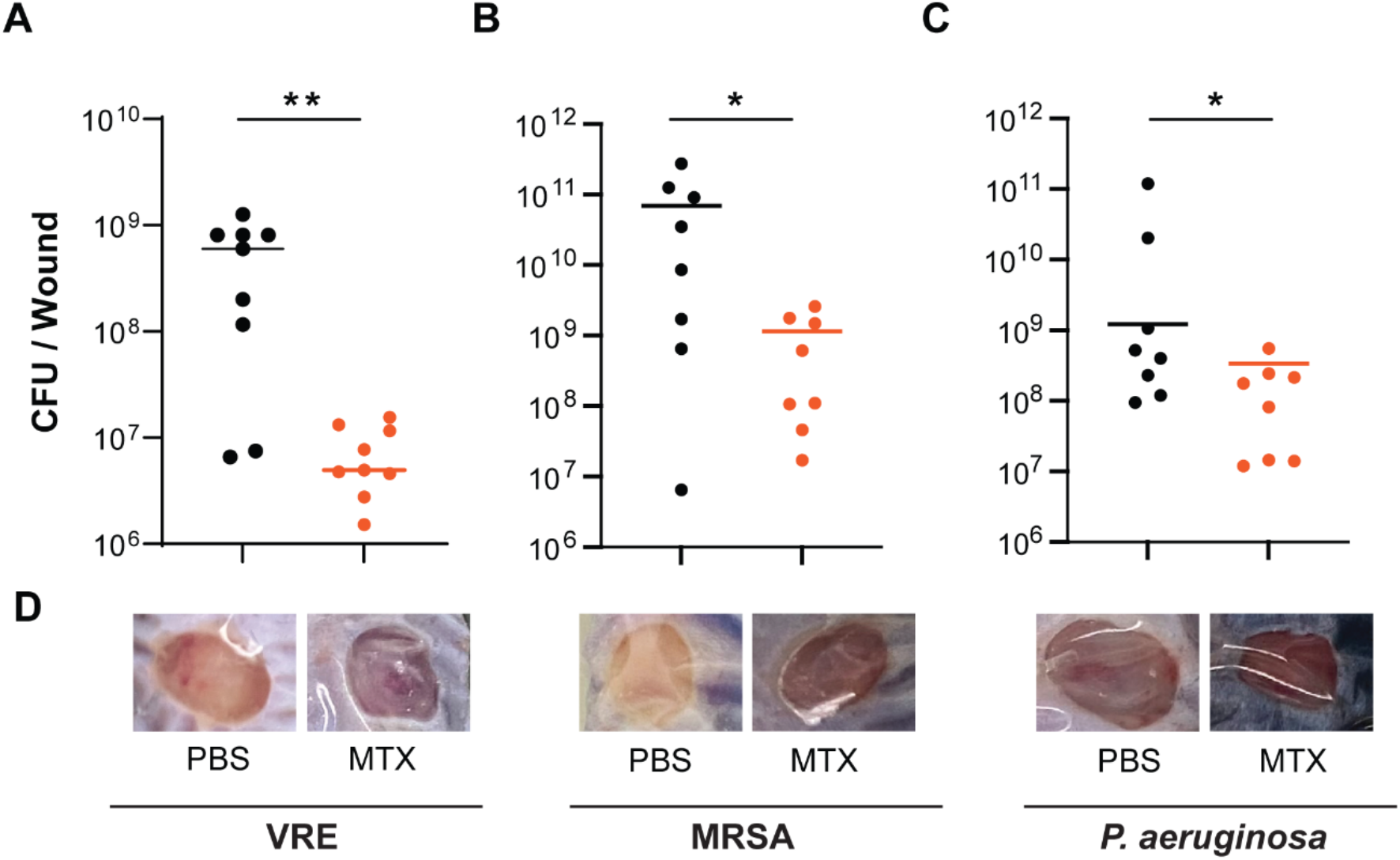
MTX exhibits potent antibiotic activity *in vivo*. **A-C.** Comparison of VRE (**A**), MRSA (**B**) and *P. aeruginosa* (**C**) CFU per infected wound treated with either PBS (black) or MTX (orange). Each symbol represents one mouse with the median indicated by the horizontal line. Data were from two independent experiments with 4-5 mice per experiment. Statistical analysis was performed using the non-parametric Mann-Whitney Test to compare ranks, *p ≤ 0.05 and **p ≤ 0.01. **D.** Representative images of VRE, MRSA and *P. aeruginosa* infected wounds treated with PBS or MTX.

### MTX synergizes with vancomycin to inhibit VRE *in vitro* and *in vivo*

We investigated whether MTX synergizes with vancomycin in inhibiting the growth of VRE using a modified MIC assay in which the concentration of MTX was kept constant at sub-MIC (0.515 μg/mL) **(Fig. 2A and fig. S2A)**, while the concentrations of vancomycin were increased by 2-fold from 0.0625 μg/mL to 75 μg/mL. As expected, VRE started to grow just below its breakpoint of 18 μg/mL vancomycin and grew without any inhibition at 1.5 μg/mL vancomycin **(Fig. 2A)**. However, the growth of VRE was potently inhibited at 0.125 μg/mL vancomycin in the presence of sub-MIC MTX **(Fig. 2A)**, indicating a 140-fold reduction of vancomycin MIC in the presence of sub-MIC MTX **(table S2)**. Combinatorial MTX and vancomycin treatment was initially bacteriostatic but became bactericidal by 24 h **(fig. S2B)**. Similarly, we tested combination of MTX (0.515 μg/mL) with other antibiotics against VRE. In the presence of MTX, the MIC for ceftriaxone, daptomycin, ciprofloxacin, chloramphenicol and Penicillin G was reduced by 8, 8, 4, 4 and 2-fold, respectively **(table S2)**, suggesting a general enhancement against VRE.

**Fig. 2.**
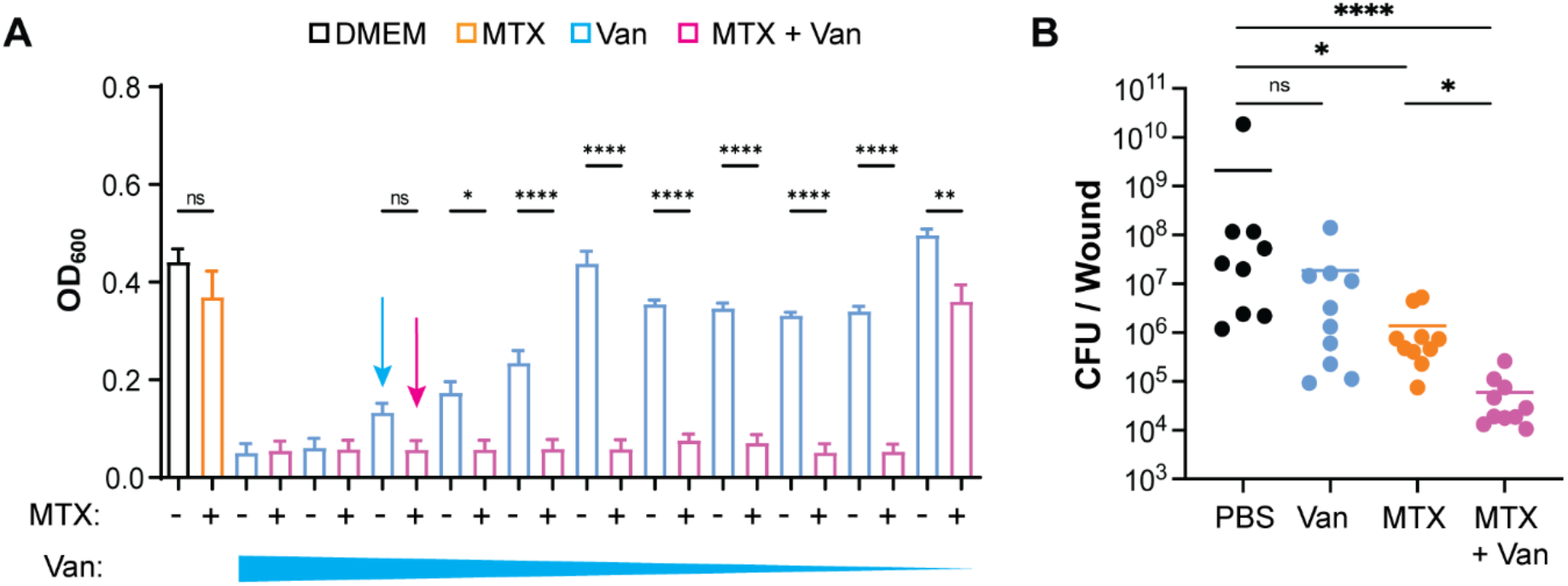
MTX and vancomycin synergize to inhibit VRE *in vitro* and *in vivo*. **A.** Comparison of VRE growth in DMEM medium (black), in the presence of MTX (0.515 μg/mL) (orange), in the presence of decreasing concentrations (75 – 0.0625 μg/mL) of vancomycin (blue), and combination of MTX and vancomycin (pink). Arrows represent breaking point where vancomycin concentration alone (blue, 18μg/mL) starts to differ from vancomycin concentration in presence of MTX (pink). Data (mean ± SEM) were derived from three independent experiments with four technical replicates for each sample per experiment. Statistical analysis was performed using ordinary one-way ANOVA, followed by Tukey’s multiple comparison test. **B.** Comparison of VRE CFU per wound without treatment (PBS), or treated with vancomycin, or MTX, or vancomycin plus MTX. Data were from two independent experiments with 4-5 mice per experiment. Each symbol represents one mouse with median indicated by the horizontal line. Statistical analysis was performed using Kruskal Wallis test with uncorrected Dunn’s post-test. For all analyses, NS denotes Non-significant, *p ≤ 0.05; **p ≤ 0.01; ***p ≤ 0.001 and ****p ≤ 0.0001.

We further tested the synergy between MTX and vancomycin in the wound infection model. Mice were first intraperitoneally injected with 100 mg/kg (human equivalent dose of ~8 mg/kg) of vancomycin, which was lower than the minimum equivalent recommended dose (>10 mg/kg) to treat Gram-positive bacterial infections in humans (*27*). Then, mice were subjected to wound infection with VRE and MTX treatment as described in **Fig. 1A**. As shown in **Fig. 2B**, vancomycin alone reduced the median CFU by 11-fold and MTX alone reduced the median CFU by 42-fold, whereas vancomycin and MTX combination reduced the median CFU by 1000-fold. Thus, MTX and vancomycin synergize to inhibit VRE both *in vitro* and *in vivo*.

### MTX kills VRE by inducing reactive oxygen species and DNA damage

MTX has been shown to stimulate the formation of reactive oxygen species (ROS) in hepatocytes (*28*). Because ROS can kill bacteria directly (*29*), we examined the role of ROS in MTX-mediated inhibition of bacterial growth by measuring the MIC of MTX under oxic and anoxic conditions. Under oxic conditions, VRE growth was completely inhibited by MTX at 1.6 μg/mL, whereas under anoxic conditions, 51.2 μg/mL of MTX was required to completely inhibit VRE growth **(Fig. 3A)**, a 32-fold decrease in MTX MIC in the presence of oxygen. Quantification of intracellular ROS production using dihydrorhodamine 123 (DHR123) revealed that the sub-MIC of MTX (0.515 μg/mL), but not vancomycin (4 μg/mL), induced significant ROS production **(Fig. 3B)**. Consistently, MTX also induced elevation of 8-hydroxy-2’-deoxyguanosine (8-OHdG), a by-product of ROS-induced DNA damage, to a level similar to that produced by 0.1 mM H_2_O_2_. **(Fig. 3C)**. Addition of vancomycin to MTX-treated cultures did not further increase the levels of ROS and 8-OHdG. Furthermore, when the ROS scavenger MitoTEMPO was added to VRE cultures, VRE growth in the presence of vancomycin plus MTX was similar to that in the presence of vancomycin alone **(Fig. 3D),** indicating that the effect of MTX in synergizing with vancomycin was completely abolished by ROS scavenger. These results suggest that induction of ROS and DNA damage is a key mechanism by which MTX exhibits antibiotic activity and synergizes with vancomycin.

**Fig. 3.**
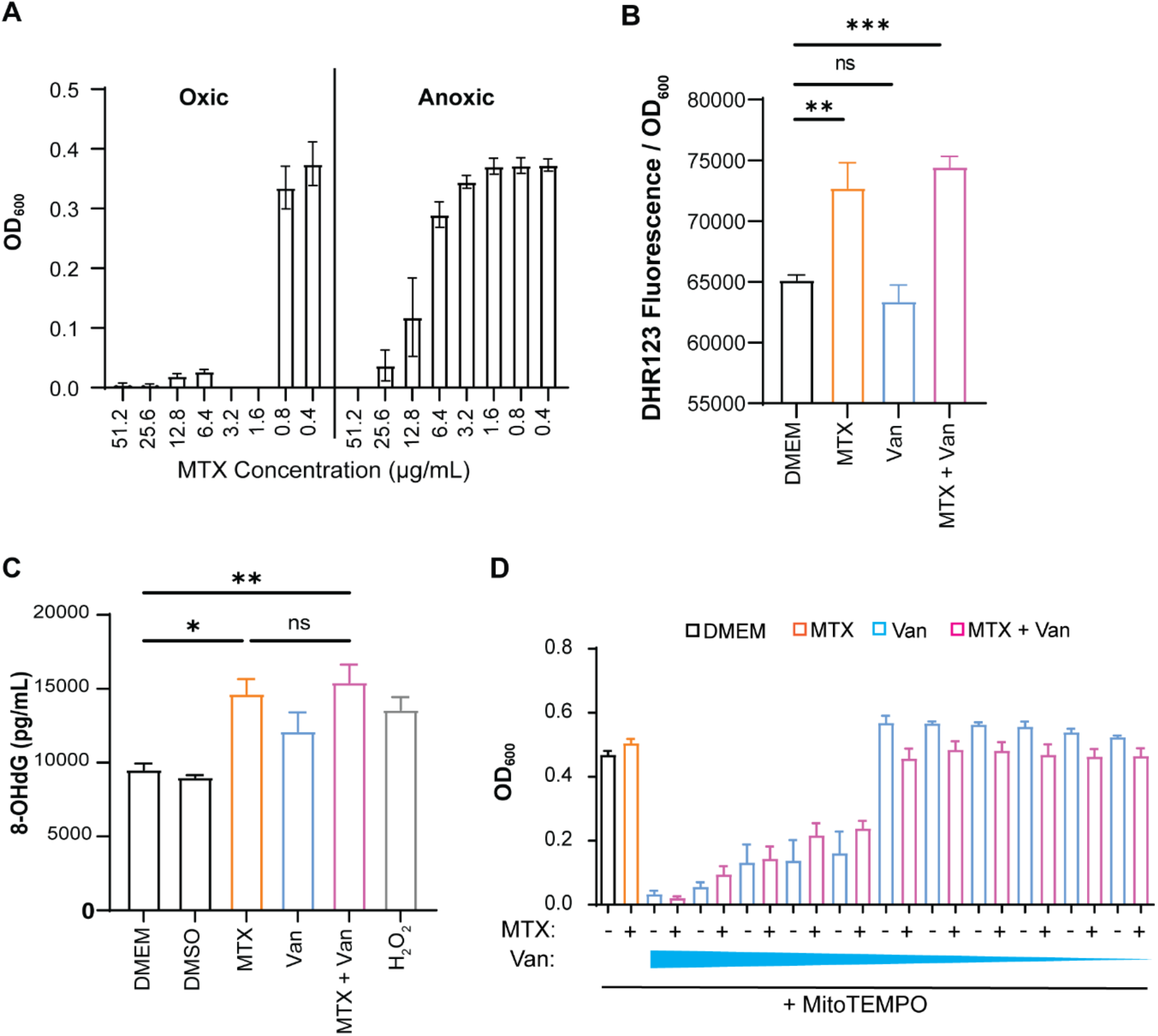
MTX induces production of reactive oxygen species and DNA damage in bacterial cells. **A.** Comparison of VRE growth (OD_600_) in oxic and anoxic conditions in the presence of decreasing concentrations of MTX. Data (mean ± SEM) are summary of three independent experiments. **B.** Comparison of intracellular ROS levels, as measured by DHR123 fluorescence, in VRE cultures treated with MTX (0.515 μg/mL), vancomycin (4 μg/mL), or both. **C.** Comparison of 8-OHdG levels, as measured by ELISA, in VRE cultures treated with MTX (0.515 μg/mL), vancomycin (4 μg/mL), or both. H_2_O_2_ (0.1 mM) was added into the VRE culture as positive control. Data (mean ± SEM) in **C** and **D** are summary from three independent experiments each. Statistical analysis was performed using ordinary one-way ANOVA, followed by Tukey’s multiple comparison test, NS p > 0.05; *p ≤ 0.05; **p ≤ 0.01; and ***p ≤ 0.001. **D.** Comparison of VRE growth in DMEM medium (black), in the presence of MTX (0.515 μg/mL) (orange), in the presence of decreasing concentrations (75 – 0.0625 μg/mL) of vancomycin (blue), and with a combination of MTX and vancomycin (pink). MitoTEMPO was added into all cultures. Data (mean ± SEM) were derived from three independent experiments for each sample per experiment.

### Vancomycin-treated VRE display increased permeability to MTX

We noticed that MTX lowered the sensitivity of VRE strains to vancomycin but did not further sensitize vancomycin-susceptible *E. faecalis* or other vancomycin-susceptible bacterial species **(Table 2)**, suggesting that the bacterial vancomycin resistance mechanism itself might play a role in the observed synergistic effect between sub-MIC doses of MTX and vancomycin. Vancomycin interferes with bacterial cell wall synthesis and therefore increases cell wall permeability (*30*). We tested if vancomycin increases the uptake of MTX by measuring MTX’s fluorescence emission at 685 nm (*31*). As shown in **Fig. 4A**, the fluorescence intensity of VRE doubled in the presence of MTX as compared to medium alone. The fluorescence intensity quadrupled when VRE were treated with both MTX and vancomycin. Consistently, mass-spectrometry quantification of intracellular MTX showed that MTX levels were ~6x higher in VRE cultures in the presence of both MTX and vancomycin than MTX alone **(Fig. 4B)**. Moreover, more propidium iodide (PI) was taken into VRE at 6 h following treatment with both MTX and vancomycin as measured by flow cytometry **(Fig. 4C)** and fluorescence microscopy **(Fig. 4D)** compared to MTX treatment alone. These results show that vancomycin increases the uptake of MTX probably due to its interference with bacterial cell wall synthesis, which is linked to the resistance mechanism itself in VRE strains, indicating that the synergy between MTX and vancomycin is due to vancomycin-induced uptake of MTX, which in turn kills bacteria by inducing ROS and DNA damage.

**Fig. 4.**
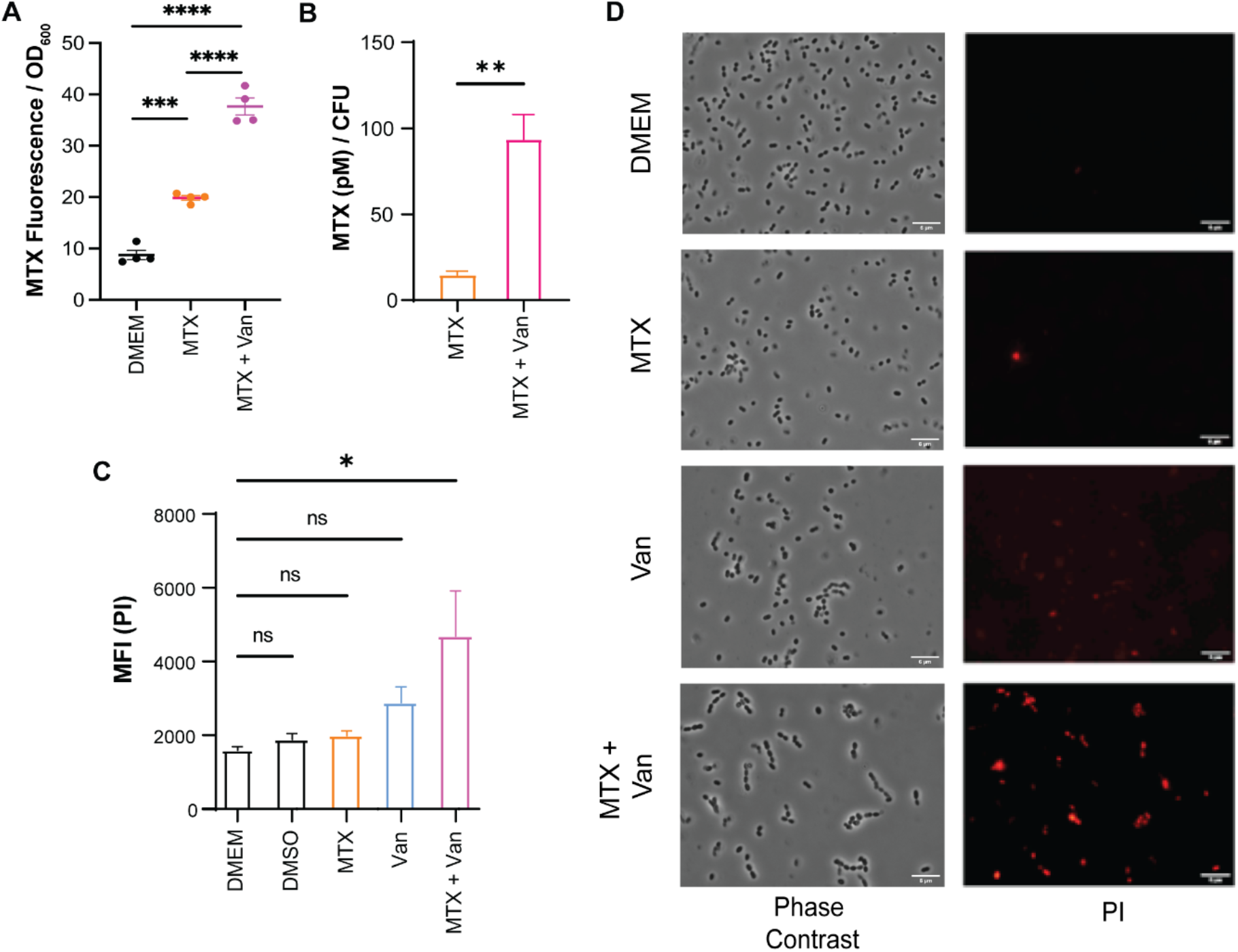
Vancomycin-treated VRE bacterial cells have increased permeability to MTX. **A.** MTX uptake by VRE after 6 h treatment with MTX (0.515 μg/mL) alone and in combination with vancomycin (4 μg/mL). Each dot represents one independent experiment. **B.** Mass-spectrometry quantification of intracellular MTX of VRE cultures treated with MTX (0.515 μg/mL) and in combination with vancomycin (4 μg/mL) for 1 h. Data (mean ± SEM) are summary of five independent replicates. Statistical analysis was performed using unpaired T-test with Welch’s corrections, ****p ≤ 0.0001. **C.** Comparison of PI uptake by VRE after 6 h treatment with MTX (0.515 μg/mL) alone, vancomycin alone (4 μg/mL), or in combination. **D.** Epifluorescence microscopy images of VRE stained with PI after 6 h treatment with MTX, vancomycin, or both. Scale bar: 5 μm. **A** and **C.** Data (mean ± SEM) are summary of at least three independent experiments. Statistical analysis was performed using ordinary one-way ANOVA, followed by Tukey’s multiple comparison test, NS p > 0.05; *p ≤ 0.05; **p ≤ 0.01; ***p ≤ 0.001 and ****p ≤ 0.0001

**Table 2.**
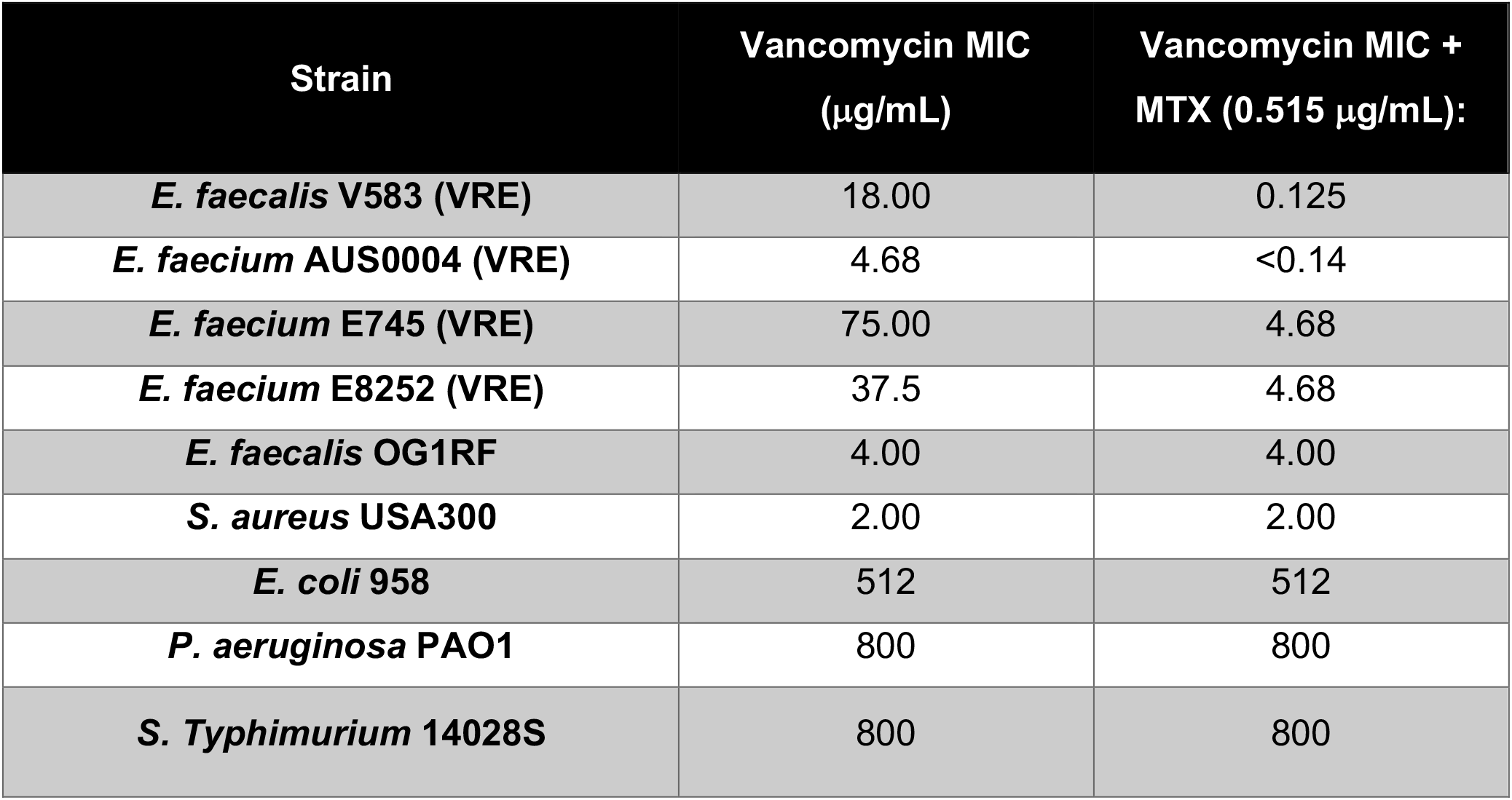
Vancomycin resistant strains have lower vancomycin MIC in presence of a sub-inhibitory dose of MTX.

### Mutation in a DEAD/DEAH box helicase confers resistance to MTX

To further investigate the mechanism by which MTX inhibits bacterial growth, we performed *in vitro* evolution to select for spontaneous mutants that were resistant to MTX. We serially passaged VRE in medium with increasing concentrations of MTX from a sub-MIC concentration of 0.515 μg/mL to 2.84 μg/mL. Among 16 colonies obtained, one exhibited the highest MTX resistance at MIC of 20 μg/mL **(Fig. 5A, table S3)** or a 12.5-fold increase over the parental VRE. This mutant, henceforth named as VRE MTX^R^, was as sensitive to vancomycin alone as the parental strain (MIC = 18 mg/mL), but was over 300-fold more resistant to vancomycin in the presence of MTX (MIC = 12.5 μg/mL). Nevertheless, VRE MTX^R^ had similar growth kinetics as the parental VRE in the absence of drugs **(Fig. 5B)**. Whole genome sequencing (WGS) revealed the presence of four mutations in the genome of VRE MTX^R^ **(Table S3)**. Among them, a point mutation (G --> A) occurred in a gene predicted to encode a DEAD/DEAH box helicase, resulting in a substitution of glycine (G) by arginine (R) at the amino acid position 389.

**Fig. 5.**
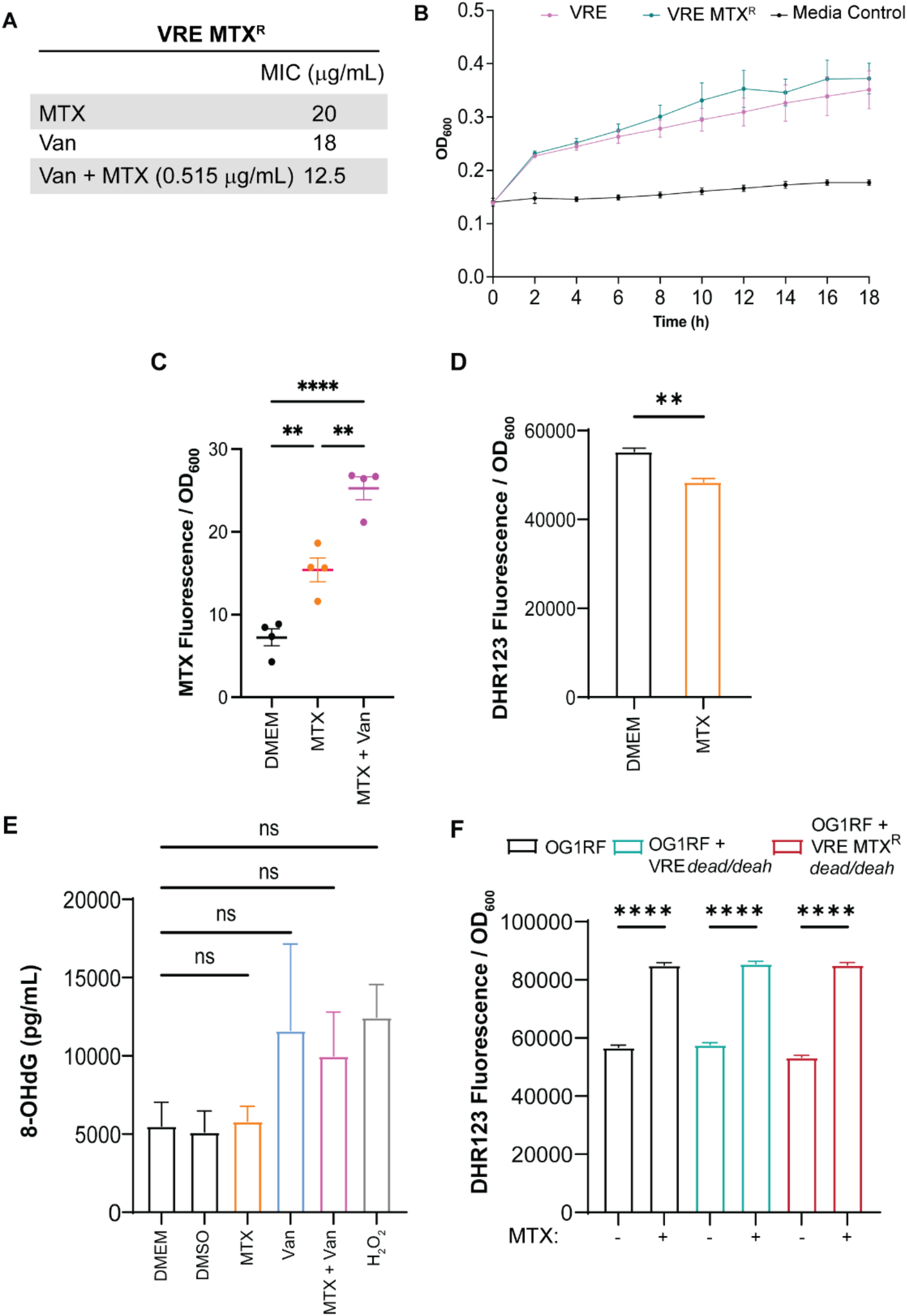
VRE MTX^R^ exhibits increased MTX uptake but not increased ROS production and DNA damage. **A.** Comparison of MIC for MTX alone, vancomycin alone, vancomycin in the presence of 0.515 mg/mL MTX between the parental VRE and VRE MTX^R^. Each dot represents one independent experiment. **B.** Growth curve of VRE and VRE MTX^R^ in DMEM**. C.** MTX uptake by VRE MTX^R^ after 6 h treatment with MTX (0.515 μg/mL) alone and in combination with vancomycin (4 μg/mL)**. D.** Analysis of ROS levels in VRE MTX^R^ treated with MTX (0.515 μg/mL). **E.** ELISA measurements of 8-OHdG levels in VRE MTX^R^ treated with MTX (0.515 μg/mL), vancomycin (4 μg/mL), separately and in combination, and the positive control H_2_O_2_ (0.1mM). **F.** Analysis of ROS levels in *E. faecalis* OG1RF and *E. faecalis* OG1RF constitutively expressing either a wildtype (VRE *dead/deah*) or the mutant DEAD/DEAH helicase (VRE MTX^R^ *dead/deah*) treated with MTX (0.515 μg/mL). **C-F.** Data (mean ± SEM) are summary of at least three independent experiments. Statistical analysis was performed using ordinary one-way ANOVA, followed by Tukey’s multiple comparison test, NS p > 0.05; *p ≤ 0.05; **p ≤ 0.01; ***p ≤ 0.001 and ****p ≤ 0.0001.

RNA helicases of the DEAD/DEAH box family have been linked to oxidative stress resistance in bacteria (*32*). Compared to the parental VRE, VRE MTX^R^ exhibited a similar increase in MTX uptake in the presence of vancomycin **(Fig. 5C)**. However, in contrast to the parental VRE, MTX treatment did not increase the intracellular ROS or 8-OHdG in VRE MTX^R^ **(Fig. 5D-E).** In fact, VRE MTX^R^ displayed a lower baseline DNA damage or DNA damage in response to H_2_O_2_ **(Fig. 5E)**, as compared to the parental VRE **(Fig. 3D)**. These results show that MTX does not induce ROS and DNA damage in the mutant VRE, likely explaining its increased resistance to MTX.

Bioinformatic analysis of the vancomycin sensitive *E. faecalis* strain OG1RF showed that it does not possess a homologue of the DEAD/DEAH box helicase gene. To determine if the DEAD/DEAH box helicase could confer OG1RF resistance to MTX, we constitutively expressed either the wildtype (WT) or the mutant DEAD/DEAH box helicase gene in OG1RF. Despite an increase in the intracellular ROS upon MTX exposure in all strains **(Fig. 5F)**, *E. faecalis* OG1RF harbouring the WT gene (VRE *dead/deah*) and the mutated copy of DEAD/DEAH box helicase gene (VRE MTX^R^ *dead/deah*) was 2- and 8-fold more resistant to MTX than the parental OG1RF strain, respectively **(Table 3)**. Thus, the DEAD/DEAH box helicase plays a critical role in protecting the VRE MTX^R^ from the effects of MTX by reducing ROS and DNA damage.

**Table 3.**
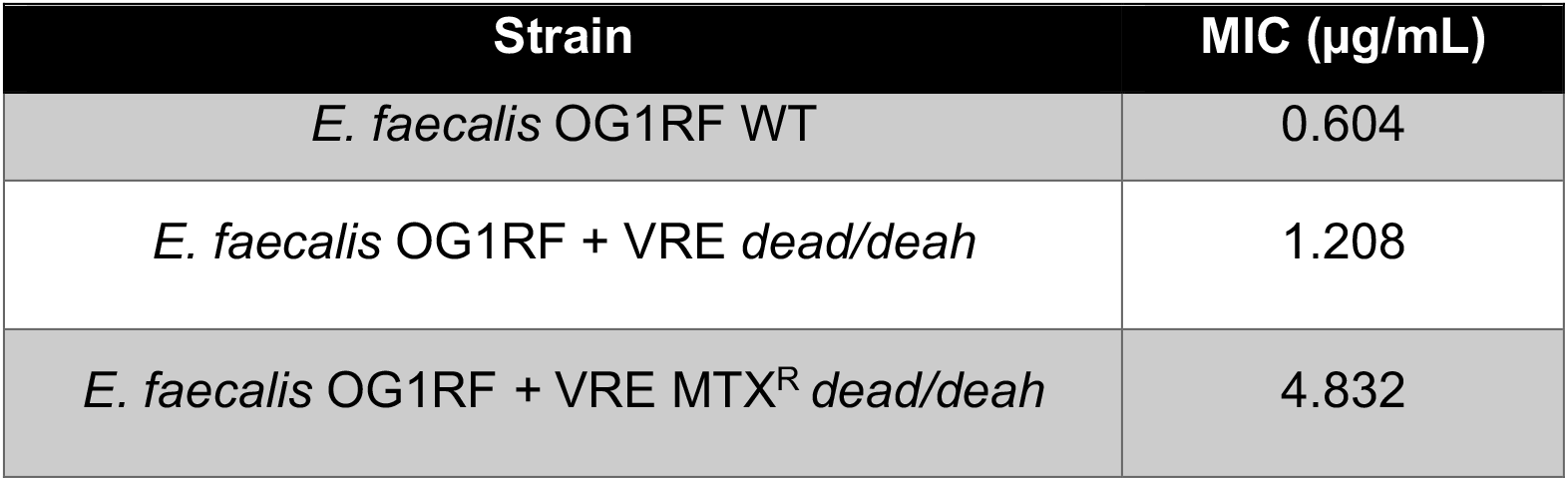
MTX MIC of *E. faecalis* OG1RF and *E. faecalis* OG1RF constitutively expressing either a wildtype (VRE *dead/deah*) or the mutant DEAD/DEAH helicase (VRE MTX^R^ *dead/deah*). MIC was established by OD_600_ to measure bacterial growth in the presence of decreasing concentrations of MTX (19.31 – 0.07 μg/mL).

### MTX enhances macrophages to eliminate bacterial infection *in vivo*

We have previously shown that MTX can reprogram macrophages toward a proinflammatory phenotype (*26*), which could lead to more effective bacterial killing. Therefore, we investigated whether MTX immunomodulatory activity contributes to reduced bacterial CFU in infected wounds *in vivo*. To distinguish between antibiotic and immunomodulatory effects of MTX, we infected wounds with VRE MTX^R^ together with MTX or PBS. MTX treatment resulted in ~290-fold fewer CFU compared to the PBS control **(Fig. 6A)**. Because the reductions in CFU were comparable (290-fold vs. 120-fold) when VRE and VRE MTX^R^ infected wounds were treated with the same amount of MTX, and because VRE MTX^R^ is resistant to MTX, the reduction of VRE MTX^R^ CFU is likely to be largely due to MTX immunomodulatory rather than antibiotic activity.

**Fig. 6.**
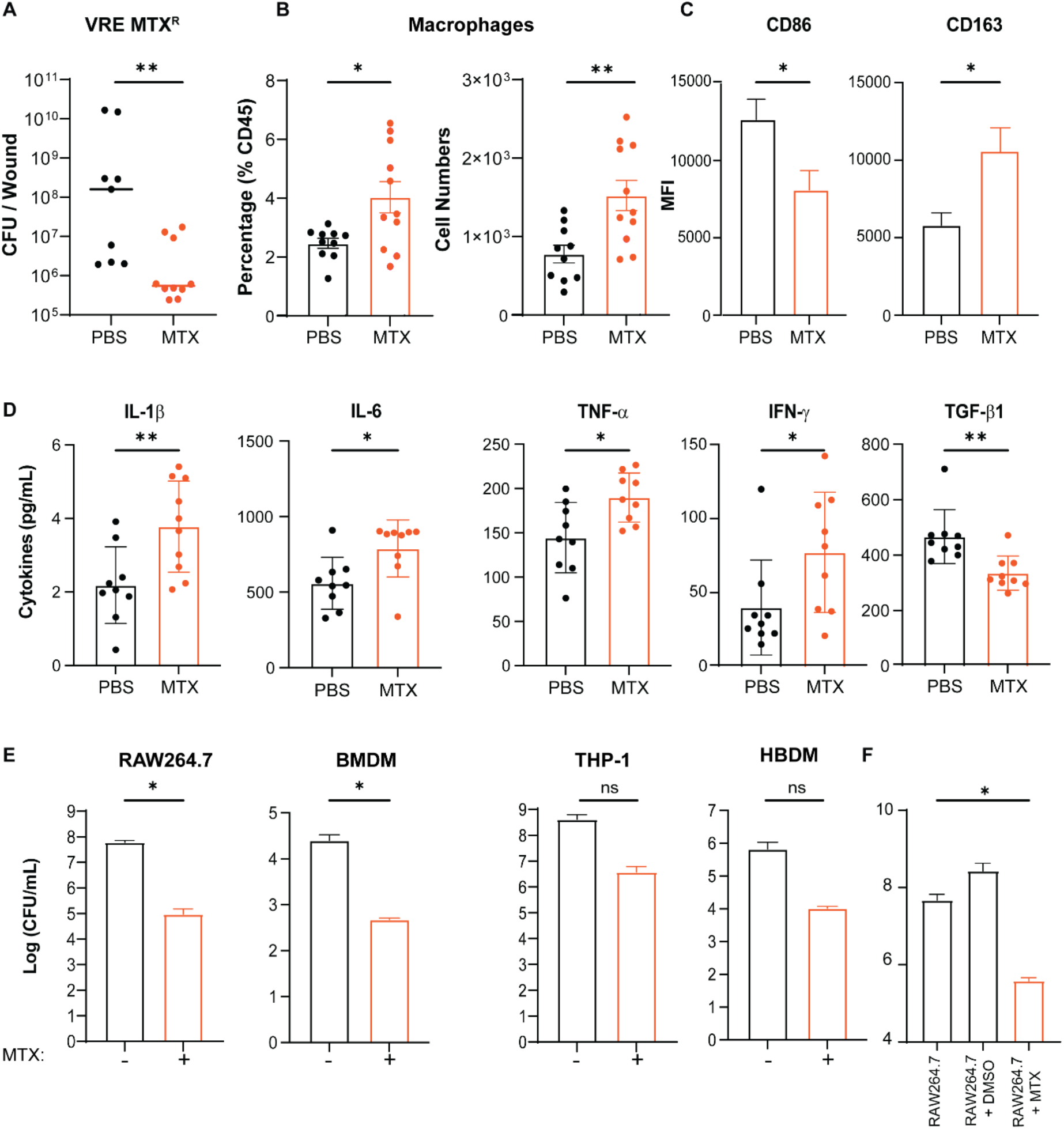
MTX treatment promotes macrophage recruitment and reprograming to a pro-inflammatory phenotype. VRE MTX^R^ infected wounds were treated for 24 h with either PBS or a single dose of MTX (10 μL of 0.515 μg/mL MTX per wound). **A.** Comparison of VRE MTX^R^ CFU in wound lysates treated with either PBS or MTX for 24 hpi. Data (mean ± median) are a summary of two independent experiments with 4-5 mice per group. **B.** Percentage and absolute numbers of macrophages recovered from infected wounds treated with PBS or MTX. Data (mean ± SEM) are summary of two independent experiments. Each dot represents one mouse. **C.** Comparison of MFI of CD86 and CD163 staining gating on CD45^+^ CD11b^+^ F4/80^+^ macrophages from infected wounds treated with PBS or MTX. Data (mean ± SEM) are from 5 mice per group. **D.** The levels of cytokines IL-1β, IL-6, TNF-α, IFN-γ and TGF-β from the lysates of infected wound treated with PBS or MTX. Data (mean ± SEM) are summary of two independent experiments. Each dot represents one mouse. **E.** Comparison of VRE CFU counts in RAW264.7, BMDM, THP-1 and HBDM in presence or absence of MTX. Data (mean ± SEM) are summary of three independent experiments. **F.** VRE CFU performed with MTX pre-treated RAW264.7 macrophage cells. Data (mean ± SEM) are summary of at least three independent experiments. Statistical analysis was performed using the non-parametric Mann-Whitney Test to compare ranks (**A**), or unpaired T-test with Welch’s corrections (**B-E**), or ordinary one-way ANOVA, followed by Tukey’s multiple comparison test (**F**). NS, not significant; p > 0.05; *p ≤ 0.05; **p ≤ 0.01; ***p ≤ 0.001 and ****p ≤ 0.0001.

To provide support for this interpretation, we quantified macrophages and neutrophils in the wounds 24 h after VRE infection and MTX treatment. VRE infected wounds treated with MTX contained twice the number of macrophages compared to that from infected wounds treated with PBS **(Fig. 6B and fig. S3A)**. Similarly, a significant increase in the percentage of neutrophils was also observed in VRE infected wounds treated with MTX **(fig. S3B)**. Flow cytometry analysis of the infection-related markers CD163, CD86, CD206 and MHCII **(fig. S3C-D)** (*33*) showed that macrophages expressed a higher level of CD163, the high affinity scavenger receptor, but a lower level of CD86 in VRE infected wounds treated with MTX as compared to PBS treatment **(Fig. 6C)**. Increased macrophage recruitment to the MTX-treated infected wounds correlated with significant increases in the levels of the pro-inflammatory cytokines 1L-1β, IL-6, TNF-α, IFN-γ in MTX-treated wounds and a lower level of the anti-inflammatory cytokine TGF-β1 **(Fig. 6D)**. Consistently, MTX treatment of infected RAW264.7 macrophages resulted in significantly higher NF-κB-driven transcription than the PBS-treated infected cells **(fig. S3E)**.

We further examined whether MTX promotes macrophage killing of intracellular bacteria. RAW264.7 macrophages and primary murine bone-marrow derived macrophages (BMDM) were infected with VRE for 3 h followed by 15 h of combined treatment with MTX and gentamicin plus penicillin, which eliminate any residual extracellular bacteria. MTX treatment resulted in ~3-log fewer intracellular CFU compared to untreated infected cells at 18 hpi **(Fig. 6E).** Similarly, MTX promoted intracellular killing of VRE in the human monocyte-like cell line (THP-1) and primary human monocyte-derived macrophages (HMDM) **(Fig. 6E)**, although the difference was not statistically significant. Moreover, MTX also enhanced macrophage killing of both Gram-positive and Gram-negative bacteria, including *E. faecium, S. aureus, P. aeruginosa*, and multi-drug resistant *E. coli* EC958 **(fig. S4)**.

To exclude a direct antibiotic effect of MTX on intracellular bacterial killing, we pre-treated RAW 264.7 macrophages with MTX overnight, washed cells to remove residual MTX in the culture, and then infected with VRE. The pre-treatment reduced intracellular CFU by ~2-log by 18 hpi **(Fig. 6F)**. Importantly, the same infection and MTX pre-treatment did not promote host cell death **(fig. S3F)** or host cell membrane permeability **(fig. S3G)**. Taken together, these results show that MTX promotes macrophage recruitment to the site of infection *in vivo* and re-programs macrophages to pro-inflammatory phenotypes to more efficiently eliminate bacteria.

### MTX enhances bactericidal activity of macrophages by stimulating lysosomal enzyme expression and activity

To determine how MTX enhances macrophage bacterial killing, we investigated whether MTX stimulates macrophage phagocytosis of bacteria. RAW264.7 macrophage cells were treated with MTX for 16 h and then incubated with fluorescent fixed VRE. After 3 h of incubation, the fluorescence of extracellular bacteria was quenched with Trypan Blue and phagocytosis of fluorescent bacteria was measured by flow cytometry and visualized by confocal microscopy. Fluorescent bacteria were visible inside RAW264.7 cells, but there was no significant difference in phagocytosis by RAW264.7 with or without MTX treatment **(fig. S5A-C)**. We also tested whether MTX stimulates macrophages to produce ROS. In the absence of infection, MTX did not induce ROS production by RAW264.7 **(fig. S5D)**. Following VRE infection, RAW264.7 cells produced ROS regardless of MTX addition. Consistently, addition of the superoxide scavenger MitoTEMPO did not reduce bactericidal activity of macrophages **(fig. S5E)**. These results suggest that neither increased phagocytosis nor ROS production explain why MTX enhances macrophage bactericidal activity.

We investigated whether MTX enhances macrophage killing of bacteria by stimulating lysosomal activity. We quantified the transcript levels of 16 lysosomal pathway genes by qRT-PCR in uninfected RAW264.7 macrophages following MTX treatment for 24 h **(fig. S6)**. Transcripts for lysosomal proteases Cathepsins D (*Ctsd*) and H (*Ctsh*), and the enzyme β-Hexosaminidase (beta subunit) (*Hexb*), which cleaves glycosides, were upregulated **(Fig. 7A and fig. S6)**. We further verified the induction of CtsD protein in MTX-treated RAW264.7 cells in both the presence and absence of bacterial infection by Western blotting **(Fig. 7B-C)**. Notably, fully processed CtsD heavy and light chains were abundant **(Fig. 7C)**, indicating activation of CtsD enzymatic activity in MTX-treated macrophages. To directly test the role of CtsD and other lysosomal proteases in bactericidal activity, we quantified the intracellular bacteria in the presence of lysosomal protease inhibitor pepstatin A. While pepstatin A alone did not inhibit macrophage killing of intracellular VRE or VRE MTX^R^, it completely abolished MTX-stimulated macrophage killing of intracellular bacteria **(Fig. 7D)**. These results show that MTX stimulates macrophages to kill bacteria by upregulating lysosomal enzyme expression and activity.

**Fig. 7.**
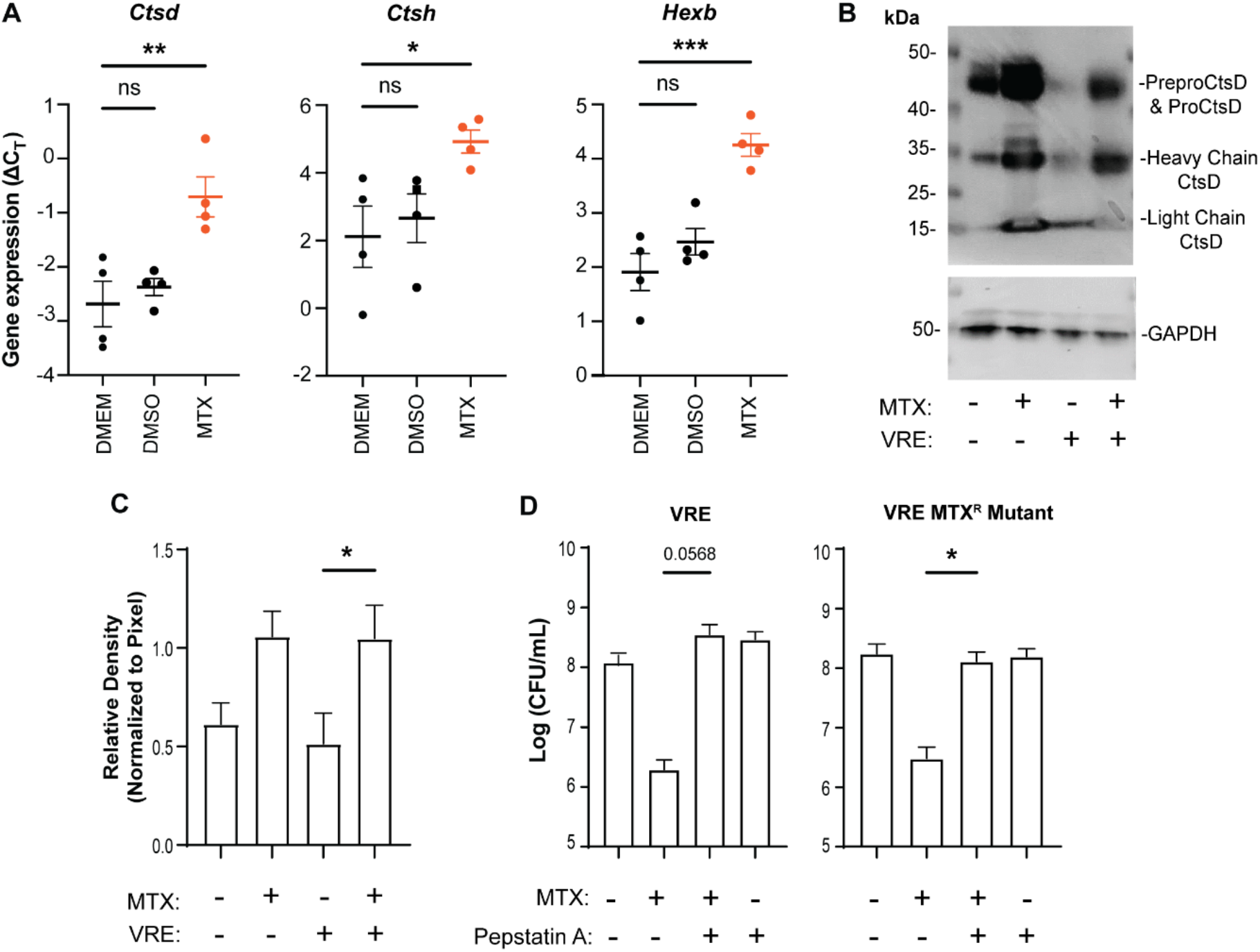
MTX enhances macrophage antimicrobial activity by stimulating lysosomal enzyme expression and activity. **A.** qRT-PCR analysis of *Ctsd, Ctsh* and *Hexb* transcript levels (ΔC_T_) in RAW264.7 cells with or without DMSO or MTX treatment overnight. Each dot represents one biological replicate. **B-C.** Western blotting analysis of whole cell lysates with anti-Cathepsin D antibody. RAW264.7 cells with (+) and without (-) VRE infection were treated with MTX (+) or left untreated (-). Whole cell lysates were separated by SDS-PAGE, transferred to membrane and probed with anti-Cathepsin D antibody or anti-GAPDH (control) (**B**). Relative band density of the CtsD heavy chain normalized to that of GAPDH (**C**). **D.** RAW264.7 cells were infected with either VRE or VRE MTX^R^ in the presence of MTX (0.515 μg/mL), pepstatin A (10 μg/mL), or both. Intracellular bacterial CFU were quantified. Data (mean ± SEM) are a summary of at least three independent experiments. Statistical analysis was performed using ordinary one-way ANOVA, followed by Tukey’s multiple comparison test, NS p > 0.05; *p ≤ 0.05; **p ≤ 0.01; ***p ≤ 0.001 and ****p ≤ 0.0001.

## Discussion

Ever-increasing antibiotic resistance requires new approaches that leverage both antibiotics and the immune system in combating bacterial infection. In this study, we investigated MTX, a chemotherapeutic initially approved for treating acute myeloid leukaemia, for its antibiotic activity and its ability to recruit and activate macrophages for bacterial clearance. We show: i) MTX possesses potent antibiotic activity against Gram-positive bacteria, ii) MTX and vancomycin synergize to overcome vancomycin-resistance, and iii) sub-MIC levels of MTX are sufficient to recruit and activate macrophages to clear bacteria in a mouse model of wound infection. We determined the molecular mechanisms that underlie MTX antibiotic activity, synergy with vancomycin, and activation of macrophage bactericidal activity.

MTX was previously reported to exhibit antibiotic activity in MIC tests performed in nutrient rich medium against *S. pneumoniae* and *S. aureus* (*34, 35*). Here we tested MTX antibiotic activity against a panel of Gram-positive and Gram-negative bacterial species and strains and found MTX to be 10-20-fold more potent against Gram-positive bacteria with MIC ~1 μg/ml. We also show that sub-MIC MTX reduced the growth of a vancomycin-resistant strain of *E. faecalis* (VRE) by ~100-fold in a murine wound infection model. To our knowledge, this is the first time MTX has been used topically to treat bacterial infections. Topical use at a low dose may limit the potential side effects of MTX administered systemically.

As a chemotherapeutic, MTX is known to induce DNA damage as well as free radical formation and lipid peroxidation in eukaryotic cells (*28*). As an antibiotic, it was previously proposed that its main mode of action was inhibition of bacterial DNA gyrase (*35, 36*). Here we show that MTX kills bacteria primarily by induction of ROS and DNA damage with three lines of evidence. First, the antibiotic activity of MTX is over 30-fold stronger in the presence of oxygen than in the absence of oxygen. Second, a sub-MIC dose of MTX is sufficient to cause significant ROS elevation and DNA damage in the bacterial cells. Third, addition of superoxide scavenger MitoTEMPO to the culture completely abolished MTX’s synergistic antibiotic activity with vancomycin. This mechanism of action is supported by the generation of an MTX-resistant mutant VRE, carrying a mutation in the gene encoding a predicted DEAD/DEAH box helicase. Enzymes of the DEAD/DEAH box family are RNA helicases implicated in many processes (*34, 43*), including oxidative stress resistance in some bacterial species (*32*). Compared to the parental VRE, VRE MTX^R^ had similar uptake of MTX, but produced lower levels of ROS, and had significantly reduced DNA damage. Furthermore, heterologous expression of the VRE wildtype and mutant DEAD/DEAH box helicase genes in *E. faecalis* OG1RF, which lacks this gene, was sufficient to confer MTX resistance by 2- and 8-fold, respectively. These results suggest that the induction of ROS and DNA damage is a key mechanism by which MTX exhibits antibiotic activity.

Clinically, combinatorial therapy using antibiotics with different mechanisms of action and different targets is used to prevent antibiotic resistance (*37*). In this study, we found that MTX and vancomycin are a highly effective combination against vancomycin resistant strains both in *in vitro* culture and in a murine model of wound infection. Interestingly, the observed synergy was limited to vancomycin-resistant strains, not vancomycin-sensitive strains, because the vancomycin resistant cells display increased permeability to MTX, thereby enabling elevated ROS production and DNA damage and therefore synergy. It is probable, that in VRE strains, vancomycin induces the resistance mechanism, leading to cell wall remodelling which in turn facilitates MTX uptake. This finding may have broader implications for the treatment of vancomycin-resistant bacterial infections and suggests the possibility of combination therapies where vancomycin may act as a permeability enhancer in strains otherwise resistant to this antibiotic.

In addition to possessing direct antibiotic activity, MTX also stimulates macrophages to more effectively clear bacteria both *in vitro* and *in vivo*. Because augmented macrophage killing activity occurred with sub-MIC concentrations of MTX, we conclude that MTX acts primarily by activating macrophages, rather than its antibiotic activity. Consistent with this interpretation, MTX-stimulated macrophage killing of bacteria is as effective for Gram-positive as for Gram-negative bacteria both *in vitro* and *in vivo*. Pre-treatment of RAW264.7 macrophages-like cells with MTX enhanced VRE killing, indicating re-programming of macrophages for enhanced bactericidal activity. Furthermore, MTX stimulated macrophages clear VRE MTX^R^ nearly as efficiently as the parental VRE strain in the wound infection, further supporting that re-programming macrophages by MTX promotes bacterial clearance.

ROS and lysosomal activity form two crucial arms of intracellular bacterial killing (*20*). *E. faecalis* can escape intracellular killing by manipulating the endosomal pathway to prevent lysosomal fusion (15). We show that MTX-treated macrophages display an enhanced bactericidal activity, rather than an enhanced phagocytosis rate, due to higher levels of expression of lysosomal enzymes including cathepsin D which has bactericidal activity against both Gram-positive and Gram-negative bacteria (*40*). Inhibition of cathepsin D using Pepstatin A ablated MTX-driven killing, whereas inhibiting ROS with MitoTEMPO had minimal effect on MTX-driven killing, suggesting that MTX-induced ROS in macrophages unlikely contributes significantly to intracellular bacterial killing. These results suggest that MTX re-programs macrophages to express an elevated levels of lysosomal enzymes and therefore stronger bactericidal activity.

In the more complex system of wound infection *in vivo*, MTX also augments macrophages to kill bacteria through additional immunoregulatory mechanisms. MTX has previously been shown to inhibit inflammatory responses and induce apoptosis in innate and adaptive immune cells (*36*) and paradoxically, to induce inflammatory responses and overexpression of M1 markers and NF-κB in macrophages in a dose-dependent manner (*25, 41*). MTX was also shown to induce inflammatory responses in adult mice following intraperitoneal injection (*41*). In our study, we showed that MTX serves as an immune attractant in infected wounds, is associated with higher levels of pro-inflammatory cytokines, and the recruited macrophages have higher levels of CD163 and lower expression of CD86. CD163, a macrophage scavenger receptor, is important for bacterial clearance, with CD163^-/-^ mice being highly susceptible to *S. aureus* infection (*42*). By contrast, CD86 can both enhance and impair immune responses to infection, likely depending on the context (*43*). Taken together, these results show that MTX promotes immunological changes within infected wounds, including recruitment of macrophages with enhanced bactericidal activity, that may contribute to better bacterial clearance.

In summary, we show that MTX possesses potent antibiotic activity by inducing ROS and DNA damage, re-sensitizes vancomycin resistant bacterial strains to vancomycin, and enhances immune cell mobilisation and intracellular clearance of bacteria. Our findings support further evaluation of MTX, especially in combination with vancomycin, for treating wound infections by vancomycin-resistant and other bacteria.

## Limitations of this study

Further validation of MTX as an antibiotic and an immunomodulatory agent, incorporating factors of mode of delivery, time of treatment and dosage will be required to confirm the suitability of this drug as a treatment for bacterial infection. We evaluated the efficacy of MTX in a murine wound excision model of VRE, MRSA and *P. aeruginosa* infection, but studies in larger animal models and ultimately humans are needed to extend these results. For simplicity, and with an eye toward uncovering mechanistic details, we focused on the effect of MTX upon innate immune cells, and in particular macrophages following a single dose regime. The effect of multiple doses of MTX, as well as the effect of MTX on other cells (both immune and otherwise) which make up the wound bed milieu are both important avenues for further investigation. Importantly, prolonged use of this compound has never been evaluated in infection contexts, but some data suggest that bacteria may be able to metabolise MTX (*69*) which could impact the spectrum of use and could also have implications in the context of polymicrobial infections, if one species metabolically depletes MTX. An important final consideration is that MTX, as a cancer therapeutic, possesses multiple mechanisms of action and potential broad off-target effects. Moreover, tissues display slow release of MTX (*36*), underscoring that the impact of MTX on tissue healing (following bacterial clearance) will need to be closely monitored. Therefore, treatment using MTX would need to be strategic, for example as adjuvant therapy or as a last resort in cases of antibiotic resistant infections, particularly vancomycin resistant bacteria.

## Materials and Methods

### Study Design

The study’s objective was to determine the efficacy and the mechanism of action of MTX as an antimicrobial and host-targeted immunotherapy for wound infections. The efficacy of MTX was investigated using a mouse wound excisional model whereby bacterial burden (CFU) was measured 24 h post-infection. Cells from *E. faecalis* V583 infected mouse wounds were profiled by flow cytometry to identify changes in cell populations between MTX versus no treatment. Cytokines levels from treated and non-treated *E. faecalis* V583 infected wounds were also investigated to investigate MTX’s mechanism of action. Additionally, *in vitro* assays with primary and cell lines of murine and human macrophages were designed to identify the individual or combined contributions of MTX and vancomycin in the enhanced bacterial killing observed *in vivo*. The group sizes for each mouse strain included at least three mice per group with at least two independent experiments to confirm results, which was sufficient to ensure statistical significance, as previously established (44, 45). Experiments were not blinded, and there were no exclusions of data or exclusion criteria to report in this study.

### Ethics statement

All animal experiments were performed with approval from the Institutional Animal Care and Use Committee (IACUC) in Nanyang Technological University, School of Biological Sciences under protocol ARF-SBS/NIE-A19061.

### Mouse wound excisional model

The procedure for mouse wound infections was modified from a previous study (15). Briefly, male C57BL/6 mice (6-8 weeks old, 22 to 25 g; NTU, Singapore) were anesthetized with 3% isoflurane. Following dorsal hair trimming, the skin was then disinfected with 70% ethanol before creating a 6-mm full-thickness wound using a biopsy punch (Integra Miltex). Bacteria (10^7^ CFU) were added to the wound site followed with addition of either 10 μL of PBS or 10 μL MTX (0.515 μg/mL). Then, the wound site was sealed with a transparent dressing (Tegaderm 3M). Where co-treatment with vancomycin was performed, intraperitoneal injections (IP) of PBS or vancomycin (100 mg/kg in a maximum volume of 100 μL) were performed before the biopsy punch. After 24 h, mice were euthanized and a 1 cm by 1 cm squared piece of skin surrounding the wound site was excised and collected in sterile PBS. Skin samples were homogenized, and the viable bacteria enumerated by plating onto BHI plates.

### Bacterial strains and growth conditions

Bacterial strains used in this study are listed in **table S4**. Bacterial strains were grown using Brain Heart Infusion (BHI) broth and agar (Becton, Dickinson and Company). For experiments performed under anoxic conditions, Oxoid AnaeroGen 3.5L sachet (Thermo Fisher Scientific) was used to create an anoxic atmosphere in the Oxoid chamber. Bacterial strains were streaked from glycerol stocks stored at −80 °C, inoculated and grown overnight statically for 16-20 h either in 10 mL of liquid BHI broth or DMEM + 10% FBS medium. Cells were harvested by centrifugation at 8000 RPM (25°C) for 5 min. The supernatant was discarded, and the pellet was then resuspended in either DMEM + 10% FBS or sterile PBS to an optical density at 600 nm (OD_600nm_) of 0.7 for VRE, equivalent to 2–3×10^8^ colony forming units (CFU).

### Antimicrobial and minimum inhibitory concentration assays

Bacterial growth assays were carried out in complete DMEM medium as described previously (46). 2 μl of overnight cultures grown in DMEM were added to 200 μl of medium in a 96-well plate with the indicated concentrations of MTX and/or vancomycin. In some assays, MitoTEMPO was also added to a final concentration of 80 μM. The OD_600_ at the zero-time point was established. Bacteria were grown statically 96-well plates at 37°C for up to 24 h. Final OD_600_ measurements were acquired using a Tecan M200 microplate reader.

### Growth curve assay and kinetic killing assay

Overnight cultures were diluted 1:100 into 96-well plates containing complete DMEM, vehicle (DMSO) or MTX (0.515 μg/ml, Sigma-Aldrich) and grown at 37°C in a Tecan M200 microplate reader. Every hour OD_600_ measurements were acquired up to 24 h. For the kinetic killing assay, overnight cultures of VRE were diluted to a starting CFU equivalent to 10^6^ CFU/mL in 50 mL tubes with 10 mL of DMEM containing MTX (0.515 μg/mL) and/or vancomycin (4 μg/ml) and grown at 37 °C. At time points up to 24 h post inoculation, 20 μl of culture was removed, serially diluted in sterile PBS, and spot plated onto BHI agar for CFU calculation.

### MTX uptake assay

VRE overnight cultures were diluted 10-fold into 10 mL of complete DMEM in 50 mL tubes containing either MTX (0.515 μg/ml), or MTX (0.515 μg/ml) and vancomycin (4 μg/ml). After 6 h, 1 mL aliquots were removed from each sample and washed 3 times with PBS. Next, 200 μL of each test sample was transferred to black-walled 96-well plates to measure MTX fluorescence (excitation = 610 nm, emission = 685 nm) by using a Tecan M200 microplate reader. MTX levels were normalized to OD_600_ to account for differences in *E. faecalis* V583 viability.

### Bacterial cells permeability assay

VRE overnight cultures were diluted 10-fold into 10 mL of complete DMEM in 50 mL tubes containing either MTX (0.515 μg/ml), vancomycin (4 μg/ml) or MTX (0.515 μg/ml) and vancomycin (4 μg/ml). After 6 h, 1 mL aliquot was removed and washed 3X with PBS prior to addition of PI (1:1000). Cells were then fixed in 4% PFA for 15 min prior to PI fluorescence analysis using a BD LSRFortessa X-20 Cell Analyzer (Becton Dickinson). In addition, epifluorescence microscopy VRE images after permeability assay were acquired using a 63× oil objective (Zeiss) fitted onto a Zeiss AxioObserver.Z1 inverted widefield microscope (Carl Zeiss, Göttingen, Germany). Acquired images were visually analyzed using using ImageJ.

### Liquid chromatography-mass spectrophotometry

Sample extraction and measurement followed the published reports with modifications (47, 48). Overnight cultures of VRE were diluted in a ratio of 1:10 in DMEM + 10% FBS and statically incubated in DMEM alone, 1 μM of MTX and/or vancomycin 4 μg/mL at 37 °C. Bacteria were harvested at 0, 1, 8 and 24 h and CFU were determined by plating serial dilutions on BHI agar medium. The cell-free supernatant was collected by filtration through a 0.22-μm filter. MTX was extracted by adding ice cold 50:50 acetonitrile/methanol. After vortexing, the mixture was centrifuged and the supernatant was collected and evaporated to dryness in a vacuum evaporator. The dry extracts were redissolved in 50:50 water/methanol for liquid chromatography-mass spectrometry (LC-MS) analysis. The calibration curve was prepared by spiking MTX standard solutions into blank medium which proceeded as described above. LC-MS analysis was performed with Agilent 1290 ultrahigh pressure liquid chromatography system coupled to an electrospray ionization with iFunnel Technology on 6490 triple quadrupole mass spectrometer. Chromatographic separation was achieved by using a Waters Atlantis T3 column with mobile phases (A) 0.1% formic acid in water and (B) 0.1% formic acid in methanol. Electrospray ionization was performed in positive ion mode and MTX was quantified in multiple reaction monitoring (MRM) mode with the transitions of *m/z* 445>88 and *m/z* 445>70. Data acquisition and processing were performed using MassHunter software (Agilent Technologies). The intracellular MTX was calculated as: [MTX]_drug-only control_ – [MTX]_filtrate_.

### DNA damage measurement

VRE overnight cultures were diluted 10-fold into 10 mL of complete DMEM in 50 mL tubes containing MTX (0.515 μg/mL), vancomycin (4 μg/mL), separately and in combination, and the positive control H_2_O_2_ (0.1mM). After 6 h, 1 mL aliquot was removed. Bacterial cells were pelleted by centrifugation (8000 RPM, 5 min) and supernatant was used to determine the levels of 8-hydroxy-2’-deoxyguanosine (8-OHdG) using a DNA Damage ELISA competitive assay as per manufacturer’s instructions.

### Bacterial reactive oxygen species quantification

This assay was adapted from (46). *E. faecalis* V583 overnight cultures were diluted 10-fold into 200 μL complete DMEM in black-walled 96-well plates containing 50 μM DHR123 (Thermo Fisher Scientific), vehicle (DMSO), MTX (0.515 μg/ml), and/or vancomycin (4 μg/ml). Plates were incubated with no shaking at 37°C for 6 h. At the end, the optical density was measured at 600 nm to determine bacterial growth and DHR123 fluorescence (excitation = 507 nm, emission = 529 nm) was measured using a Tecan M200 microplate reader to determine cellular ROS levels. ROS levels were normalized to OD_600_ to account for differences in VRE viability.

### *In vitro* evolution of *E. faecalis* to MTX resistance

The protocol was adapted from a previously published *in vitro* evolution experiment done in *E. faecalis* V583 (*49*). Sixteen starting colonies were picked from an overnight grown plate and grown in DMEM + 10% FBS overnight for performing parallel lines of evolution experiment. 10X dilutions of overnight bacterial cultures of each strain were made in DMEM + 10% FBS containing MTX from a starting sub-MIC concentration of 0.258 μg/mL and incubated at 37°C at static conditions for 22 to 26 h. Cultures of every evolution line were examined for visible bacterial growth. Bacterial cultures were then diluted again 10X into fresh MTX-containing medium at higher concentration. This was repeated till MTX of 2.84 μg/mL was achieved. Bacterial cultures were then further passaged 2 more times in the highest concentration, prior to genomic DNA extraction using Wizard Genomic DNA Purification Kit (Promega, USA). Each of the 16 bacterial cultures were also further evaluated for MIC method, as described above, to validate their susceptibility to higher concentrations of MTX. Evolved strains were also subjected to whole genome sequencing. Raw reads were imported into CLC Genomics Workbench 8.0 (Qiagen), followed by quality trimming to remove bad quality reads. The trimmed reads were then mapped to the reference genome before the Basic Variant Detection module was used to detect for mutations using the default parameters.

### Strain construction

To construct the V583DEAD (*dead/deah* gene WT copy) and MTX^R^V583DEAD (*dead/deah* mutated gene copy) complementation plasmid, primers 1 and 2 **(table S4)** were designed with *XhoI* restriction sites. These primers flank the gene of interest and were used to amplify the DNA sequence from the isolated genomic DNA of the WT and MTX^R^ strains. In-Fusion cloning (TaKaRa Bio) was performed using primers 1 and 2 with at least 15 bp complementary sequence for ligation into vector pGCP123 (*50*), which was also digested with the same restriction enzyme. The pGCP123::V583DEAD and pGCP123::MTX^R^V583DEAD plasmid was generated in *E. coli* DH5α, verified by sequencing, and transformed into *E. faecalis* as described previously (*50*).

### Flow cytometry

Flow cytometry was performed as described in (*14*) with some modifications. Excised skin samples were placed in 1.5 ml Eppendorf tubes containing 2.5 U/ml liberase prepared in DMEM with 500 μg/ml of gentamicin and penicillin G (Sigma-Aldrich). The mixture was then transferred into 6-well plates and incubated for 1 h at 37°C in a 5% CO_2_ humidified atmosphere with constant agitation. Dissociated cells were then passed through a 70 μm cell strainer to remove undigested tissues and spun down at 1350 RPM for 5 min at 4°C. The enzymatic solution was then aspirated, and cells were blocked in 500 μl of FACS buffer (2% FBS and 0.2 mM ethylenediaminetetraacetic acid (EDTA) in PBS (Gibco, Thermo Fisher Scientific)). 10^7^ cells per sample were then incubated with 10 μl of Fc-blocker (anti-CD16/CD32 antibody, Biolegend) for 30 min, followed by incubation with an anti-mouse CD45, CD11b, and Ly6G (neutrophils), or CD45, CD11b and F4/80 (macrophages) plus CD86 and MHCII or CD163 and CD206 markers conjugated antibodies (Biolegend) (1:100 dilution) for 30 min at room temperature. Cells were then centrifuged at 500 x g for 5 min at 4 °C and washed in FACS buffer. Cells were fixed in 4 % PFA for 15 min at 4 °C, before final wash in FACS buffer and final resuspension in this buffer. Following which, cells were analysed using a BD LSRFortessa X-20 Cell Analyzer (Becton Dickinson). Compensation was done using AbC Total Antibody Compensation Bead Kit (Thermo Fisher Scientific) as per manufacturer’s instructions. To evaluate MTX cytotoxicity in RAW264.7 cells, cells were analysed using a BD LSRFortessa X-20 Cell Analyzer after being stained with PI (1:1000) for 30 min on ice.

### Cytokine analysis

Homogenized wound samples were stored at −80 °C until use. Samples were thawed on ice, and centrifuged for 5 min at 500 x g to remove cell debris. Supernatants were used to perform ELISA to quantify the levels of IL-1β (Thermo Fisher Scientific), IL-6 (Biolegend), TNF-α (Biolegend), IFN-γ (Biolegend), TGF-β1 (Thermo Fisher Scientific), according to the manufacturer’s instructions.

### Human blood derived macrophages (HBDM) and cell lines

Isolated peripheral blood (PB) primary human monocytes were purchased from StemCells Technologies. For *in vitro* differentiation of monocytes into human macrophages, isolated monocytes were cultured in complete RMPI1640 supplemented with 10% heat-inactivated fetal bovine serum (FBS) (PAA, GE Healthcare), 2 mM L-glutamine (Corning) and 1% PenStrep solution (Gibco, Thermo Fisher Scientific) in the presence of 50 ng/mL recombinant human M-CSF (Biolegend) for 7 days. The RAW264.7 murine macrophage-like cell line (InvivoGen), and the THP-1 monocytic cells derived from an acute monocytic leukemia patient cell line (ATCC), were cultured at 37°C in a 5% CO_2_ humidified atmosphere. All cells were grown and maintained in Dulbecco’s modified Eagle’s medium (DMEM) (Gibco; Thermo Fisher Scientific) with 10% heat-inactivated FBS (PAA, GE Healthcare), and 100 U of penicillin–streptomycin (Gibco, Thermo Fisher Scientific). The culture medium was replaced every three days, and upon reaching 80% confluency, cultures were passaged. RAW264.7 cells passaging was achieved by gentle cell scraping and seeding cells at a density of 3×10^6^ cells/T75 flask (Nunc; Thermo Fisher Scientific).

### Mouse bone marrow-derived macrophages (BMDM)

BMDMs were prepared as described previously (*51*). Briefly, fresh bone marrow cells were isolated from mice, plated in complete RPMI with 50 ng/mL recombinant M-CSF (Biolegend) and cultured for 6 days with medium change every 3 days.

### Intracellular infection assay

Intracellular infection assays were performed as described in (14) with some modifications. Cells were seeded at a density of 10^6^ cells/well or 8×10^5^ cells/well in a 6-well or 96-well tissue culture plate (Nunc; Thermo Fisher Scientific), respectively, and allowed to attach overnight at 37°C in a 5% CO_2_ humidified atmosphere. Cells were infected at a multiplicity of infection (MOI) of 10 for up to 3 h. Following infection, the media was aspirated, and the cells were washed three times in PBS and incubated with 150 μg/ml of gentamicin (Sigma-Aldrich) and 50 μg/ml penicillin G (Sigma-Aldrich) to kill extracellular bacteria and MTX (0.515 μg/ml), or varying concentrations of vancomycin (0.06-75 μg/ml) (Sigma-Aldrich) and MTX (0.515 μg/ml), in complete DMEM for 18-24 h to selectively kill extracellular bacteria. The antibiotic containing medium was then removed, the cells were washed 3 times in PBS before addition of 2% Triton X–100 (Sigma-Aldrich) PBS solution to lyse the cells for enumeration of the intracellular bacteria. Variations of this assay included pre-treatment of mammalian cells, prior to bacterial infection, with MTX (0.515 μg/ml) followed by antibiotic treatment only, or co-treatment of cells at the time of infection with either MitoTEMPO (80 μM) (Sigma-Aldrich) or Pepstatin A (10 μg/mL) (Sigma-Aldrich).

### NF-κB reporter assay

This assay was performed as described in (*13*) using RAW-blue cells (InvivoGen). Post treatment of RAW267.4 cells for 16 h with MTX (0.515 μg/ml) or LPS (100 ng/mL) and IFN-γ (50 ng/mL) or IL-4 (10 ng/mL) and IL-13 (10 ng/mL), 20 μl of supernatant was added to 180 μl of Quanti-Blue reagent (Invivogen) and incubated overnight at 37°C. SEAP levels were determined at 640 nm by using a Tecan M200 microplate reader.

### LDH cell viability assay

As described before (*13*), post intracellular infection assays, culture supernatants were collected from each well to measure lactate dehydrogenase (LDH) release by using an LDH cytotoxicity assay (Clontech) according to the manufacturer’s instructions. Background LDH activity was determined using mock (PBS)-treated RAW264.7 cells. Maximal LDH activity was determined by lysing cells with 1% Triton X. The percentage of cytotoxicity was calculated as follows: % cytotoxicity = [(sample absorbance – background absorbance)/(maximal absorbance – background absorbance)] × 100.

### Mammalian cell reactive oxygen species quantification

Mammalian cells were seeded at a density of 8×10^5^ in a 96-well tissue culture plate (Black Nunc; Thermo Fisher Scientific), respectively and allowed to attach overnight at 37°C in a 5% CO_2_ humidified atmosphere. Fluorescein (Abcam) was added to each well to a final concentration of (120 nM) followed by addition of the positive control (H_2_O_2_, 1mM), vehicle (DMSO), and MTX (0.515 μg/ml). Cells were then either infected at a MOI 10 for up to 6 h or left uninfected. Plates were incubated with no shaking at 37°C. At the end, the fluorescence (excitation = 490 nm, emission = 525 nm) was measured using a Tecan M200 microplate reader to determine cellular ROS levels.

### RNA isolation and qRT-PCR

To quantify the levels of RNA transcripts, total RNA was extracted from non-treated, DMSO-treated or MTX-treated RAW264.7 cells with RNeasy MinElute Kit (Qiagen) and reverse transcribed using SuperScript III First-Strand Synthesis SuperMix (Thermo Fisher), followed by amplification with KAPA SYBR Fast (Kapa Biosystems) with specific primers (Table S4) and detected by Step One Plus Real time PCR machine (Applied Biosystems). Relative quantification of gene expression was performed using the comparative Ct Method (52) where the Ct values were normalized with housekeeping gene GAPDH for comparison.

### Immunoblotting

Whole cell (WC) lysates were prepared by adding 488 μl of RIPA buffer (50 mM Tris-HCl, pH 8.0; 1% Triton X–100; 0.5% Sodium deoxycholate; 0.1% SDS; 150 mM NaCl) to the wells after intracellular infection assays, where cells were scraped and kept in RIPA buffer for 30 min at 4 °C. Prior to the addition of 74.5 μl of 1 M DTT and 187.5 μl NuPAGE LDS Sample Buffer (4X) (Thermo Fisher Scientific), cells were further mechanically disrupted by passing the lysate through a 26g size needle. Samples were then heated to 95°C for 5 min. 15 μl of cell lysate proteins were then separated in a 4–12% (w/v) NuPAGE Bis-Tris protein gel and transferred to PVDF membranes. Membranes were incubated with Tris-buffered saline, TBS (50 mM Tris, 150 mM NaCl, pH 7.5) containing 0.1% (v/v) Tween-20 (TBST) and 5% (w/v) BSA for 1 h at room temperature. Membranes were incubated with 1:1000 for rabbit α-cathepsin D (Cell Signaling Technology), or 1:1000 for rabbit α-GADPH (Cell Signaling Technology) in TBST containing 1% (w/v) BSA overnight at 4°C. Membranes were washed for 60 min with TBST at room temperature and then incubated for 2 h at room temperature with goat anti-rabbit (H+L) HRP-linked secondary antibody (Invitrogen) respectively. After incubation, membranes were washed with TBST for 30 min and specific protein bands were detected by chemiluminescence using SuperSignal West Femto maximum sensitivity substrate (Thermo Fisher Scientific). Band intensities were quantified relatively to the lane’s loading control using ImageJ (53).

### Phagocytosis Assay

*E. faecalis* V583 cells were fixed with 4% PFA for 15 min and washed thrice with PBS, prior to labelling with the membrane permeant DNA dye - Syto9 (Thermo Fisher Scientific). Bacterial cells were then washed thrice with PBS and resuspended in DMEM + 10% FBS. RAW264.7 cells were infected with MOI10 of Syto9-labelled bacterial cells and incubated for 1 hour at 37°C and 5% CO_2_. Following supernatant removal, infected cells were harvested and resuspended in PBS. The fluorescence of bacteria either free in the medium or attached to the RAW264.7 cell membranes were quenched with a final concentration of 0.01% trypan blue. As trypan blue cannot enter viable eukaryotic cells, the unquenched fluorescence reflected the bacterial cells that were internalized in viable RAW264.7 cells. After staining, cells were immediately run through the flow cytometer. All data were collected using the BD LSRFortessa X-20 Cell Analyzer and analyzed with FlowJo V10.8.1 (BD Biosciences, USA). The samples were initially gated side scatter area (SSC-A) by forward scatter area (FSC-A) to select the RAW264.7 populations. The RAW264.7 population was subsequently gated forward scatter width (FSC-W) by side scatter area (SSC-A) to remove doublet populations. The resulting singlet cell population was then assessed for Syto9 fluorescent marker.

### Fluorescence Staining

RAW264.7 cells were seeded at 2×10^5^ cells/well in a 24-well plate with 10 mm coverslips, and allowed to attach overnight at 37°C and 5% CO_2_. Infection with Syto9 fluorescent-labelled *E. faecalis* V583 was performed with MOI10 for 1 h. The coverslips seeded with cells were then fixed with 4% PFA at 4°C for 15 min, permeabilized with 0.1% Triton X–100 for 15 min at room temperature and washed thrice in PBS. Cells were then blocked with PBS supplemented with 0.1% saponin and 2% bovine serum albumin (BSA). For actin labelling, the phalloidin–Alexa Fluor 555 conjugate (Thermo Fisher Scientific, USA) was diluted 1:40 in PBS and incubated for 1 h. Coverslips were then washed 3 times in PBS with 0.1% saponin. They were then subjected to a final wash with PBS, thrice. Finally, the coverslips were mounted with SlowFade Diamond Antifade (Thermo Fisher Scientific) and sealed. Confocal images were then acquired on a 63x/NA1.4, Plan Apochromat oil objective fitted onto an Elyra PS.1 with LSM 780 confocal unit (Carl Zeiss), using the Zeiss Zen Black 2012 FP2 software suite. Laser power and gain were kept constant between experiments. Z-stacked images were processed using Zen 2.1 (Carl Zeiss). Acquired images were visually analyzed using ImageJ (53).

### Statistical analysis

Statistical analysis was done using Prism 9.2.0 (Graphpad, San Diego, CA). We used non-parametric Mann-Whitney Test to compare ranks, and one-way analysis of variance (ANOVA) with appropriate post-tests, as indicated in the figure legend for each figure, to analyze experimental data comprising 3 independent biological replicates, where each data point is typically the average of a minimum 2 technical replicates (unless otherwise noted). In all cases, a p value of ≤0.05 was considered statistically significant.

## Data and materials availability

All data associated with this study are present in the paper or the Supplementary Materials.

## Supplementary Figures and Tables

**Fig. S1.**
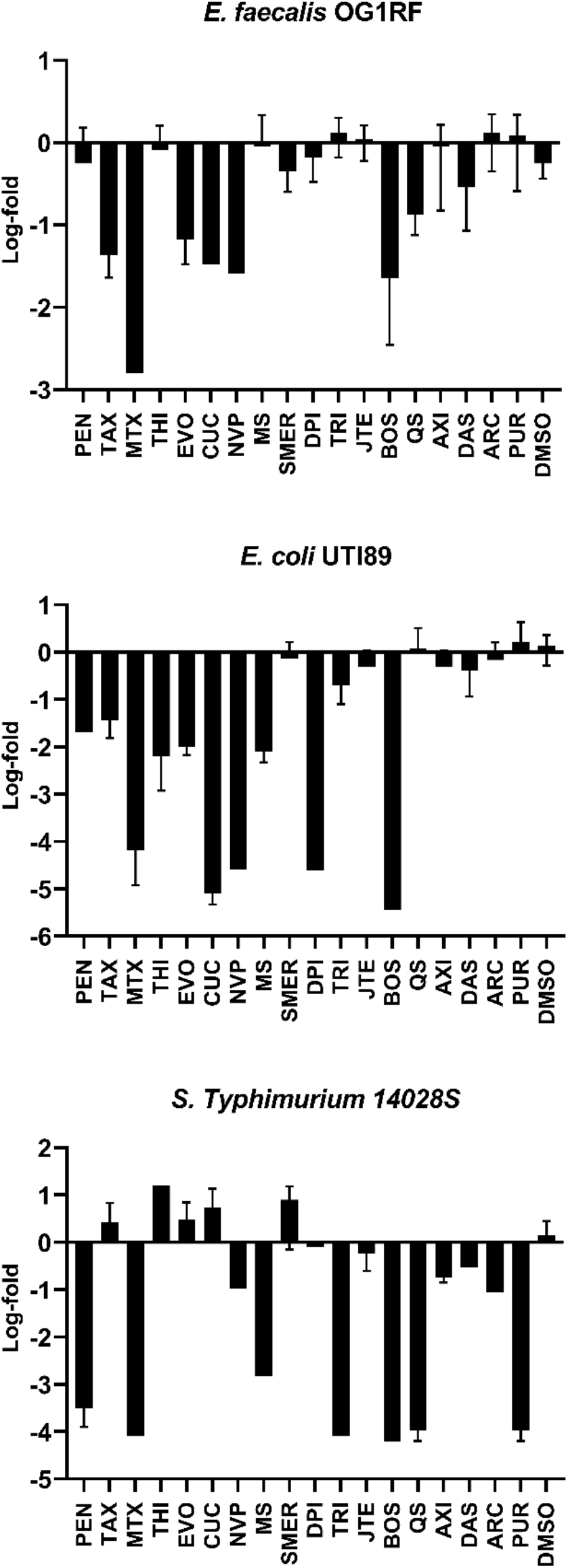
MTX induces intracellular killing of bacteria. Raw264.7 cells were infected with the indicated bacterial species for 3 h, followed by 1 h of antibiotic treatment to kill extracellular bacteria. After removal of the first antibiotic medium, and wash, a new solution with antibiotics was added together with 18 different compounds. Viable intracellular CFU were enumerated after 18 h incubation. The log-fold change was calculated with reference to the intracellular CFU of infected cells that were not treated with any compound. MTX-treated cells had lower intracellular CFU regardless of the bacterial species tested.

**Fig. S2.**
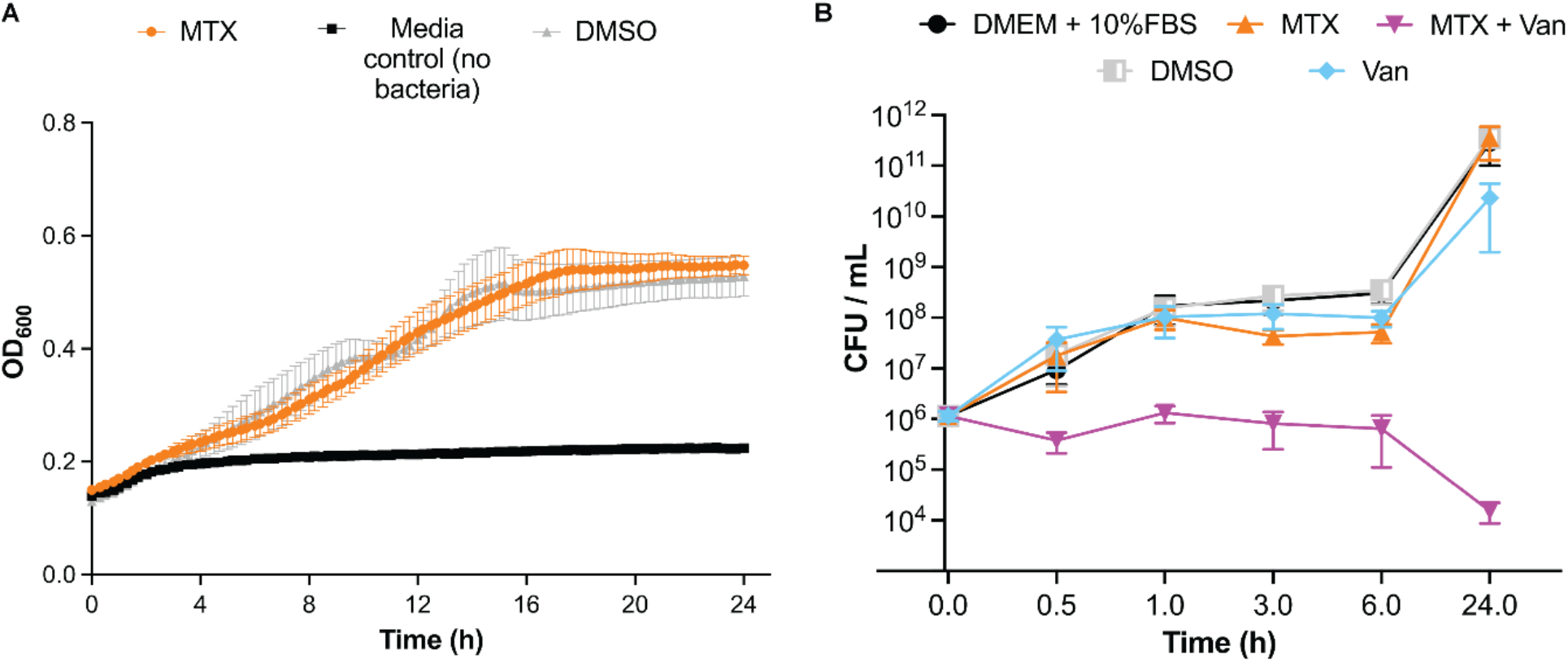
Low dose of MTX in presence of low dose of vancomycin inhibits VRE growth. **A.** Growth curve of VRE in presence of MTX (0.515 μg/mL) in DMEM for 24 h**. B.** VRE growth curve by CFU counting. In a 10 mL tube, a bacterial starting culture of 10^6^ CFU/mL was incubated with MTX, Vancomycin, separately or in combination, and vehicle (DMSO). At different time points, serial dilutions were performed in a 96-well plate and spotted on a BHI plate to establish the CFU/mL. **A and B**. Data (mean ± SEM) are a combination of at least three independent experiments.

**Fig. S3.**
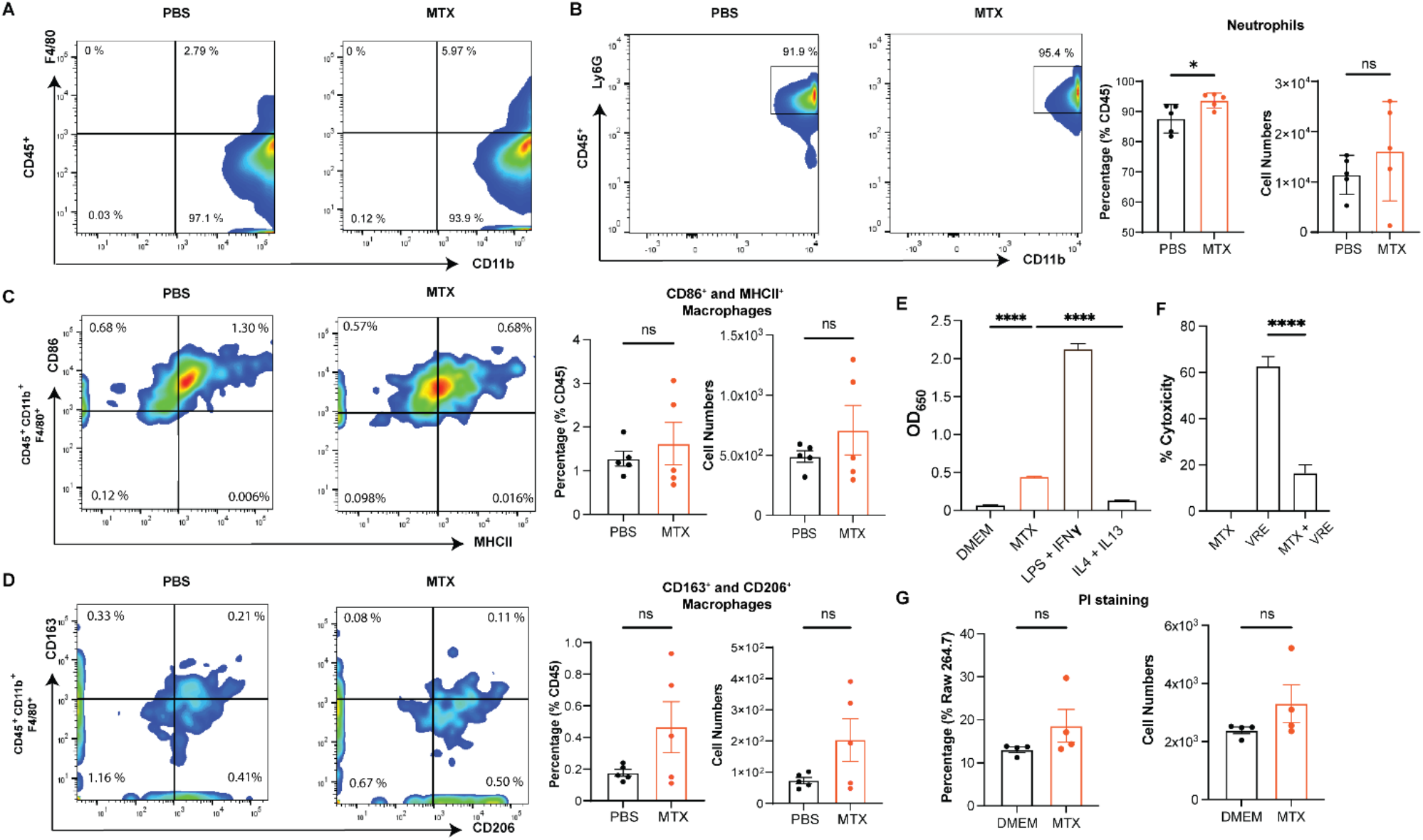
MTX treatment does not affect macrophage polarization but influences NF-κB activity. VRE Infected mouse wounds were treated for 24 h with either PBS or a single dose of MTX (0.515 μg/mL, 10 μL on the wound). **A.** Representative flow cytometry of macrophages (CD45^+^ CD11b^+^ F4/80^+^) from infected wounds treated with PBS or MTX. The number indicate percentages of cells within the gated areas. **B.** Representative flow cytometry of neutrophils (CD45^+^ CD11b^+^ Ly6G^+^) from infected wounds treated with PBS or MTX. The number indicate percentages of cells within the gated areas. Percentage and absolute numbers of neutrophils recovered from infected wounds treated with PBS or MTX. Each dot represents one mouse. **C.** Representative flow cytometry of macrophages (CD45^+^ CD11b^+^ F4/80^+^ CD86^+^ MHCII^+^) from infected wounds treated with PBS or MTX. The number indicate percentages of cells within the gated areas. Percentage and absolute numbers of CD86^+^ MHCII^+^ macrophages recovered from infected wounds treated with PBS or MTX. Each dot represents one mouse. **D.** Representative flow cytometry of macrophages (CD45^+^ CD11b^+^ F4/80^+^ CD163^+^ CD206^+^) from infected wounds treated with PBS or MTX. The number indicate percentages of cells within the gated areas. Percentage and absolute numbers of CD163^+^ CD206^+^ macrophages recovered from infected wounds treated with PBS or MTX. Each dot represents one mouse. **B-D**. Statistical analysis was performed using unpaired T-test with Welch’s corrections, NS p > 0.05. **E.** NF-κB-driven SEAP reporter activity. RAW267.4 macrophages were untreated or treated with MTX or LPS (100 ng/mL) and IFN-γ (50 ng/mL) or IL-4 (10 ng/mL) and IL-13 (10 ng/mL) 16 h prior to measurement of NF-κB-driven SEAP reporter activity. **F.** LDH activity after intracellular infection assay with MTX. RAW267.4 macrophages were infected or not with VRE MOI 10 for 3 h and then left untreated or treated with MTX 16 h in presence of antibiotic treatment prior to measurement cytotoxicity (LDH activity). While supernatant of untreated cells were considered the background noise and were subtracted from the test values, cells lysed with Triton X-100 were considered 100 %. **E-F.** Data (mean ± SEM) are a summary of at least three independent experiments. Statistical analysis was performed using ordinary one-way ANOVA, followed by Tukey’s multiple comparison test, NS p > 0.05; *p ≤ 0.05; **p ≤ 0.01; ***p ≤ 0.001 and ****p ≤ 0.0001. **G.** Percentage and absolute numbers of PI staining of RAW267.4 macrophages untreated or treated with MTX for 16 h prior to measurement. Data (mean ± SEM) are a summary of at least three independent experiments. Statistical analysis was performed using unpaired T-test with Welch’s corrections, NS p > 0.05.

**Fig. S4.**
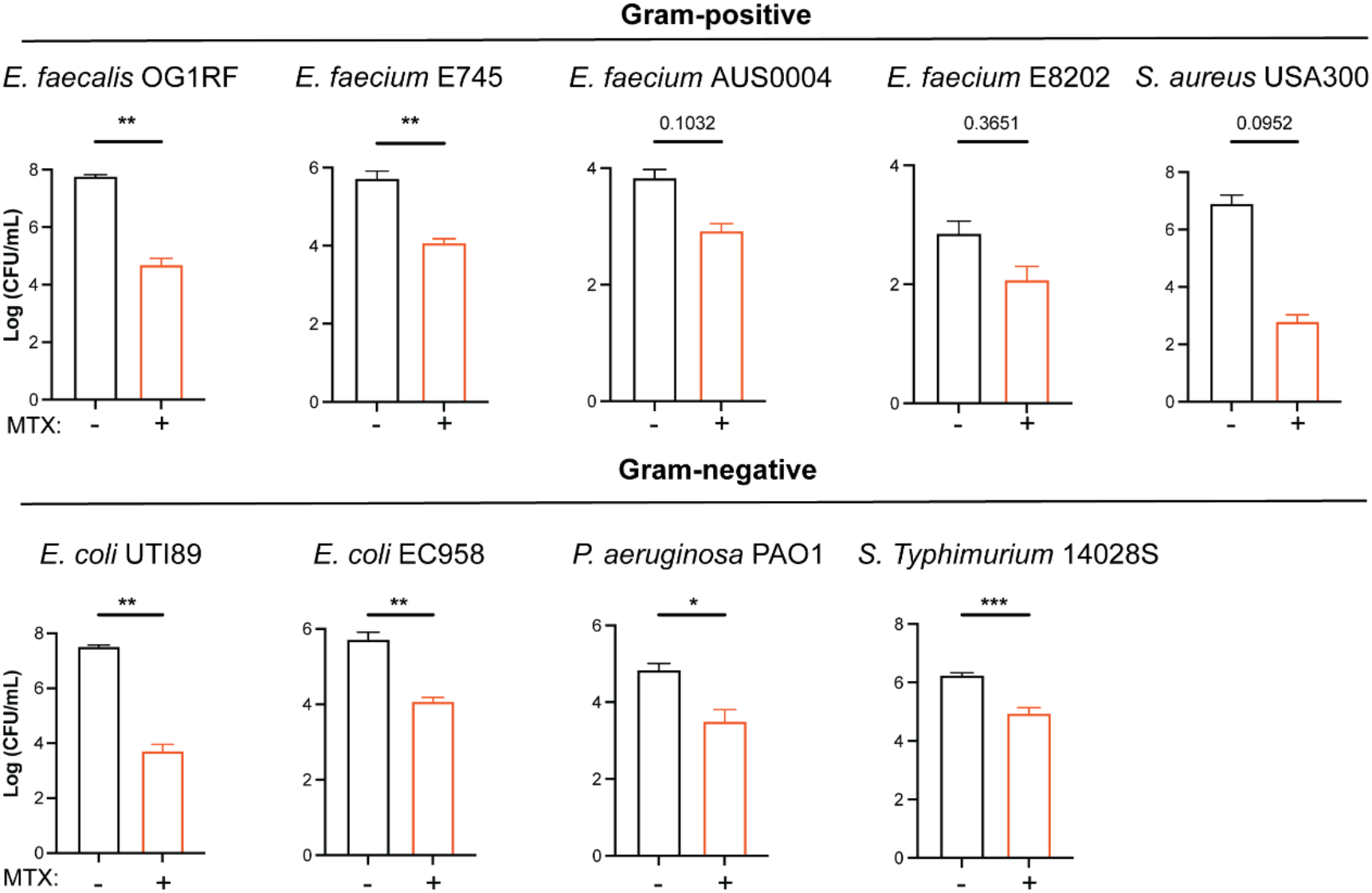
MTX as antimicrobial against multiple bacterial species *in vitro*. **A.** Comparison of VRE CFU counts of different Gram-positive and Gram-negative bacterial species in RAW264.7 in presence or absence of MTX. Data (mean ± SEM) are summary of three independent experiments. Statistical analysis was performed using the non-parametric Mann-Whitney Test to compare ranks, NS p value is shown; *p ≤ 0.05; **p ≤ 0.01; ***p ≤ 0.001.

**Fig. S5.**
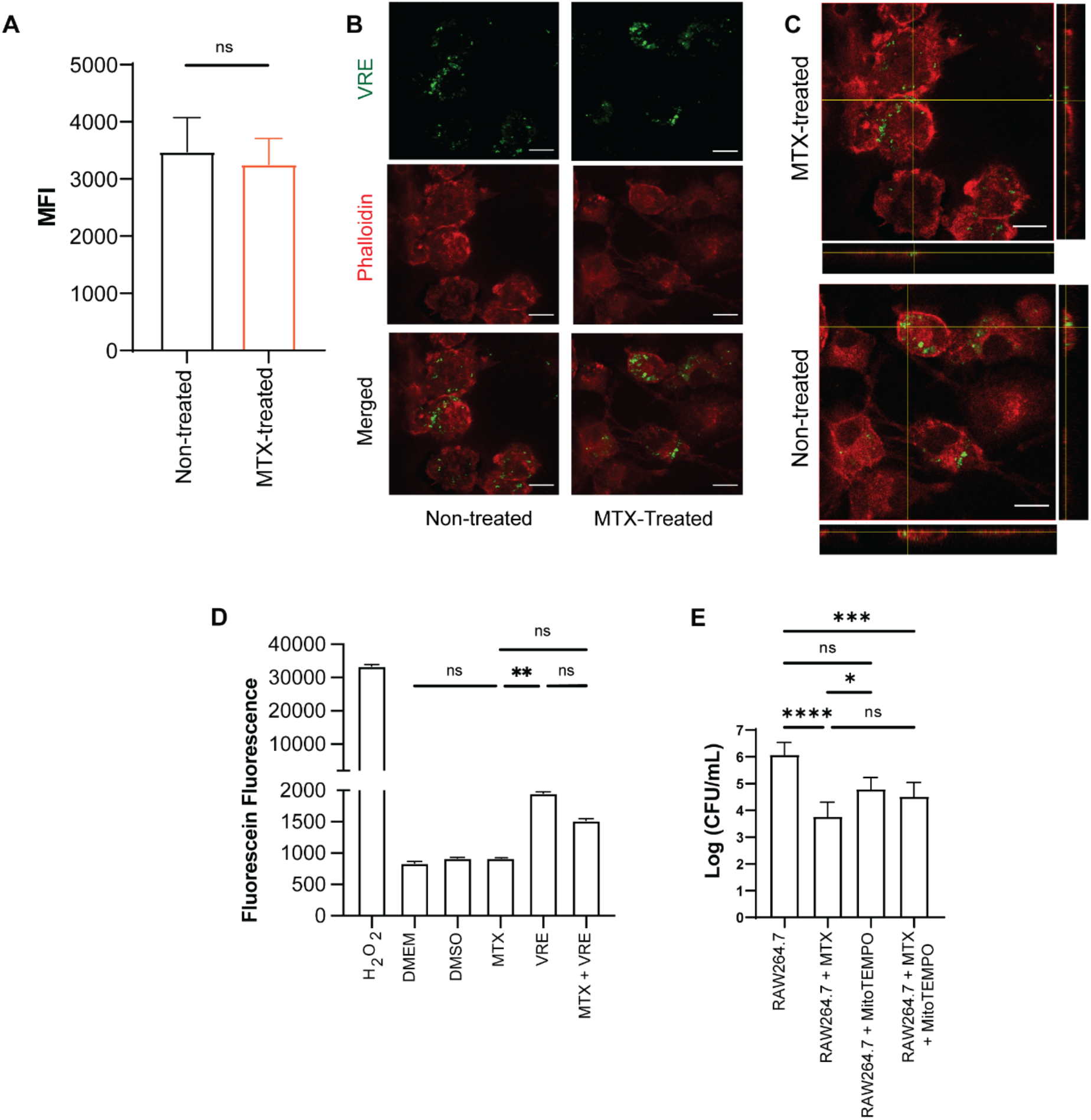
MTX does not induce phagocytosis nor ROS in RAW264.7. **A - C.** Phagocytosis assay. MTX pre-treated RAW264.7 macrophages were infected for 1 h with SYTO9-labelled VRE, quenched with Trypan Blue and then run through a flow cytometer for green fluorescence measurement. **A.** Mean Fluorescence Intensity (MFI). Data (mean ± SEM) are a summary of at least three independent experiments. Statistical analysis was performed using unpaired T-test with Welch’s corrections, NS p > 0.05. **B and C.** Representative CLSM images and orthogonal views of matching samples that were also stained with phalloidin ultra-cellular structure visualization. Scale bar: 20 μm. **D.** ROS levels of RAW264.7 macrophages untreated (DMEM) or treated with H_2_O_2_ (1 mM, positive control), DMSO, or MTX and infected or not with VRE for 6h. **E.** Comparison of VRE CFU counts in RAW264.7 in presence or absence of MTX and/or MitoTEMPO (80 μM). **D-E.** Data (mean ± SEM) are a summary of at least three independent experiments. Statistical analysis was performed using ordinary one-way ANOVA, followed by Tukey’s multiple comparison test, NS p > 0.05; *p ≤ 0.05; **p ≤ 0.01; ***p ≤ 0.001 and ****p ≤ 0.0001.

**Fig. S6.**
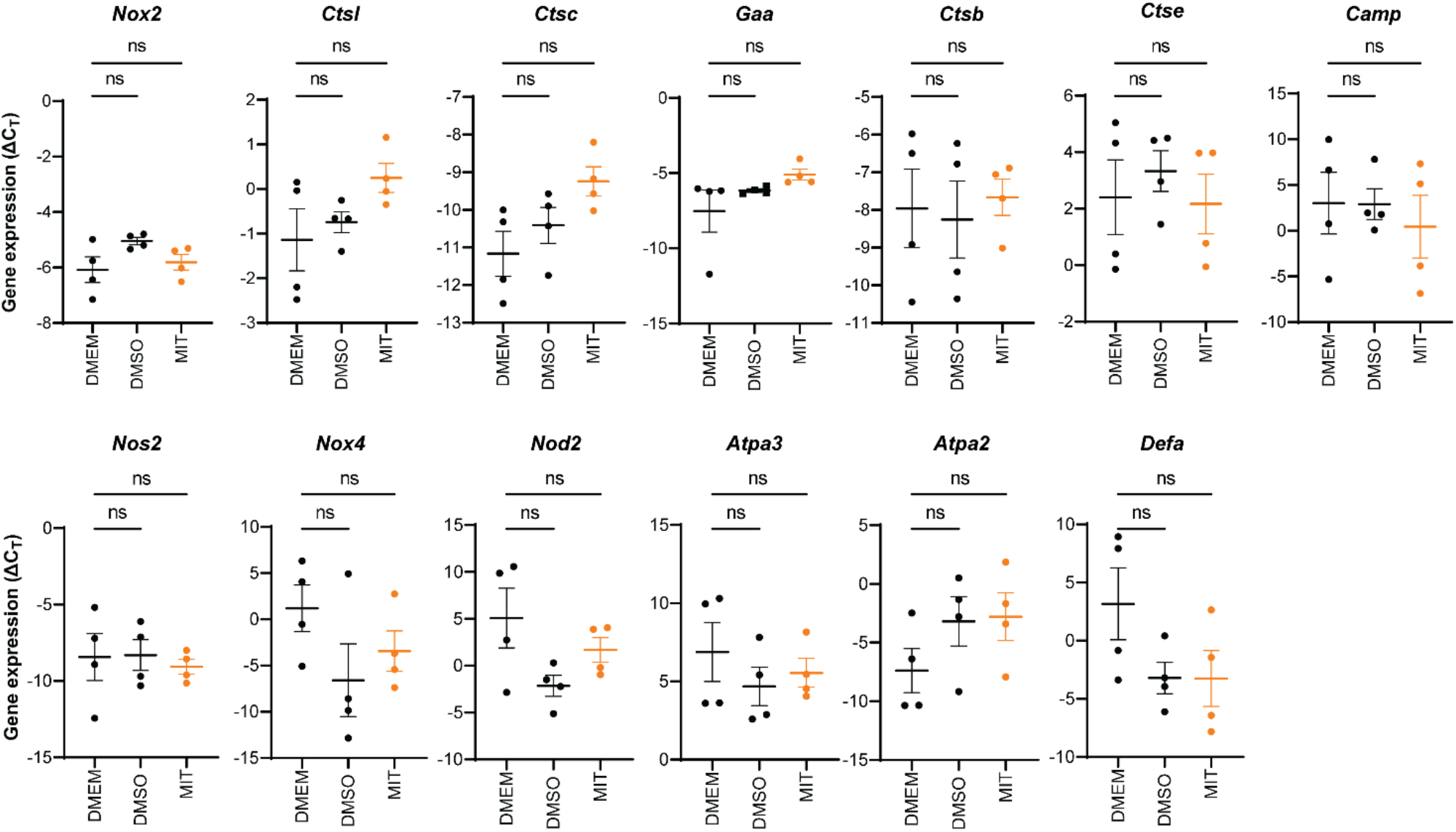
Lysosomal genes expression not affected by MTX treatment. qRT-PCR analysis of lysosomal genes transcript levels (ΔC_T_) in RAW264.7 cells with or without DMSO or MTX treatment overnight. Each dot represents one biological replicate. Statistical analysis was performed using ordinary one-way ANOVA, followed by Tukey’s multiple comparison test, NS p > 0.05.

**Table S1.** List of compounds used in initial screen.

| Abbreviation | Name | Brand |
| --- | --- | --- |
| PEN | Penfluridol | Sigma-Aldrich |
| TAX | Taxol / Paclitaxel | Sigma-Aldrich |
| MTX | Mitoxantrone HCL | Sigma-Aldrich |
| THI | Thiostrepton | Sigma-Aldrich |
| EVO | Evodiamine | Sigma-Aldrich |
| CUC | Cucurbitacin I | Sigma-Aldrich |
| NVP | NVP 231 | Sigma-Aldrich |
| MS | MS 275 | SingLab |
| SMER | SMER3 | SingLab |
| DPI | Diphenyleneiodonium Chloride | SingLab |
| TRI | Triptolide | Sigma-Aldrich |
| JTE | JTE 3 | Sigma-Aldrich |
| BOS | Bosutinib | Sigma-Aldrich |
| QS | QS 11 | SingLab |
| AXI | Axitinib | Sigma-Aldrich |
| DAS | Dasatinib | Sigma-Aldrich |
| ARC | Arcyriaflavin A | SingLab |
| PUR | Purmorphamine | Sigma-Aldrich |

**Table S2.**
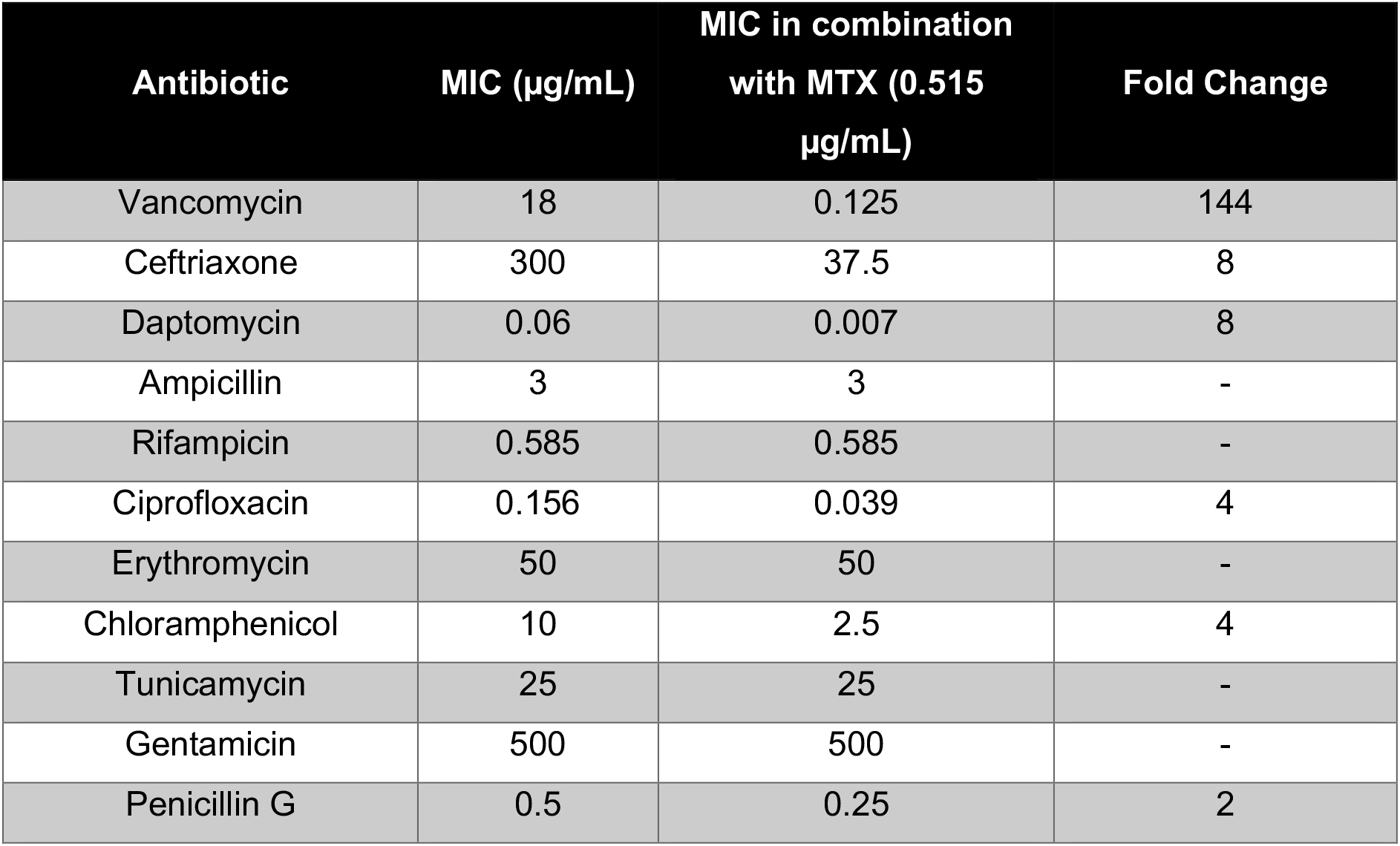
Antibiotic MIC alone or in the presence of a sub-inhibitory concentration of MTX (0.515 μg/mL).

**Table S3.** Evolution experiment strain information. Parental and VRE MTX^R^ strains MIC’s are shown.

| Strain | Information |  |  |  | Vancomycin<br>MIC (µg/mL) | Vancomycin<br>MIC + MTX<br>(0.515<br>µg/mL): | MTX<br>MIC |
| --- | --- | --- | --- | --- | --- | --- | --- |
| *E. faecalis*<br>V583 | Parental |  |  |  | 18.00 | 0.125 | 1.615 |
| *E. faecalis*<br>V583<br>MTX<sup>R</sup> | **Mutation**<br>1: G→A | **Annotation and position**<br>G389R (GGG→AGG) | **Gene**<br>DEAD/DEAH box<br>helicase | **Locus Tag**<br>EF_RS01660 | 18.00 | 12.50 | 20 |
|  | 2: G→A | intergenic (-54/+67) | Sugar<br>O-acetyltransferase | EF_RS05155 |  |  |  |
|  | 3: 2 bp→AC | insertion (805-806/912 nt) | Ethanolamine<br>ammonia-lyase subunit<br>EutC | EF_RS07835 |  |  |  |
|  | 4: (C)5→4 | insertion (171/390 nt) | Conjugal transfer<br>protein | EF_RS09020 |  |  |  |

**Table S4.**
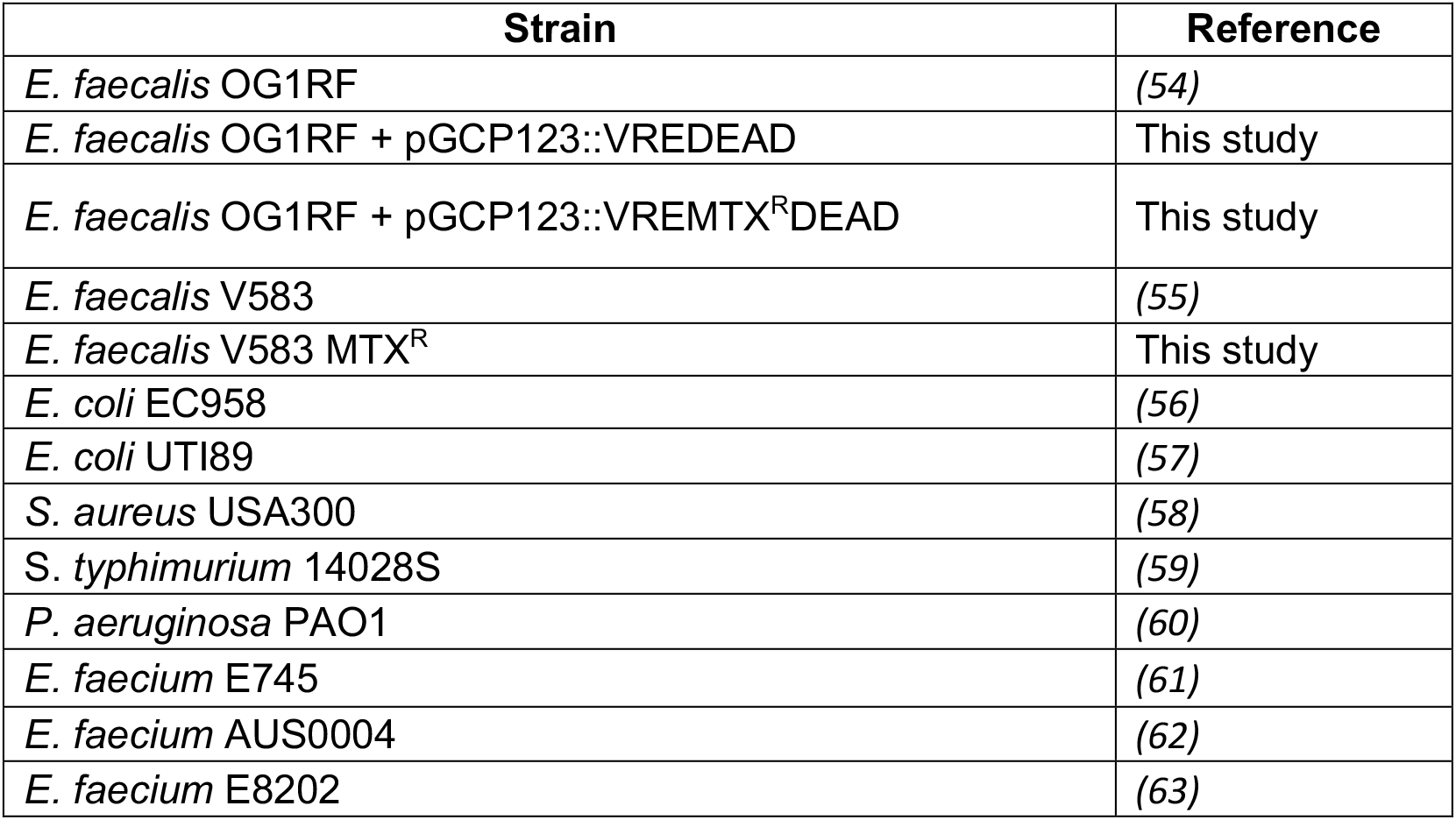
Strains used in this study.

| Strain | Reference |
| --- | --- |
| *E. faecalis* OG1RF | (54) |
| *E. faecalis* OG1RF + pGCP123::VREDEAD | This study |
| *E. faecalis* OG1RF + pGCP123::VREMTX<sup>R</sup>DEAD | This study |
| *E. faecalis* V583 | (55) |
| *E. faecalis* V583 MTX<sup>R</sup> | This study |
| *E. coli* EC958 | (56) |
| *E. coli* UTI89 | (57) |
| *S. aureus* USA300 | (58) |
| *S. typhimurium* 14028S | (59) |
| *P. aeruginosa* PAO1 | (60) |
| *E. faecium* E745 | (61) |
| *E. faecium* AUS0004 | (62) |
| *E. faecium* E8202 | (63) |

**Table S5.**
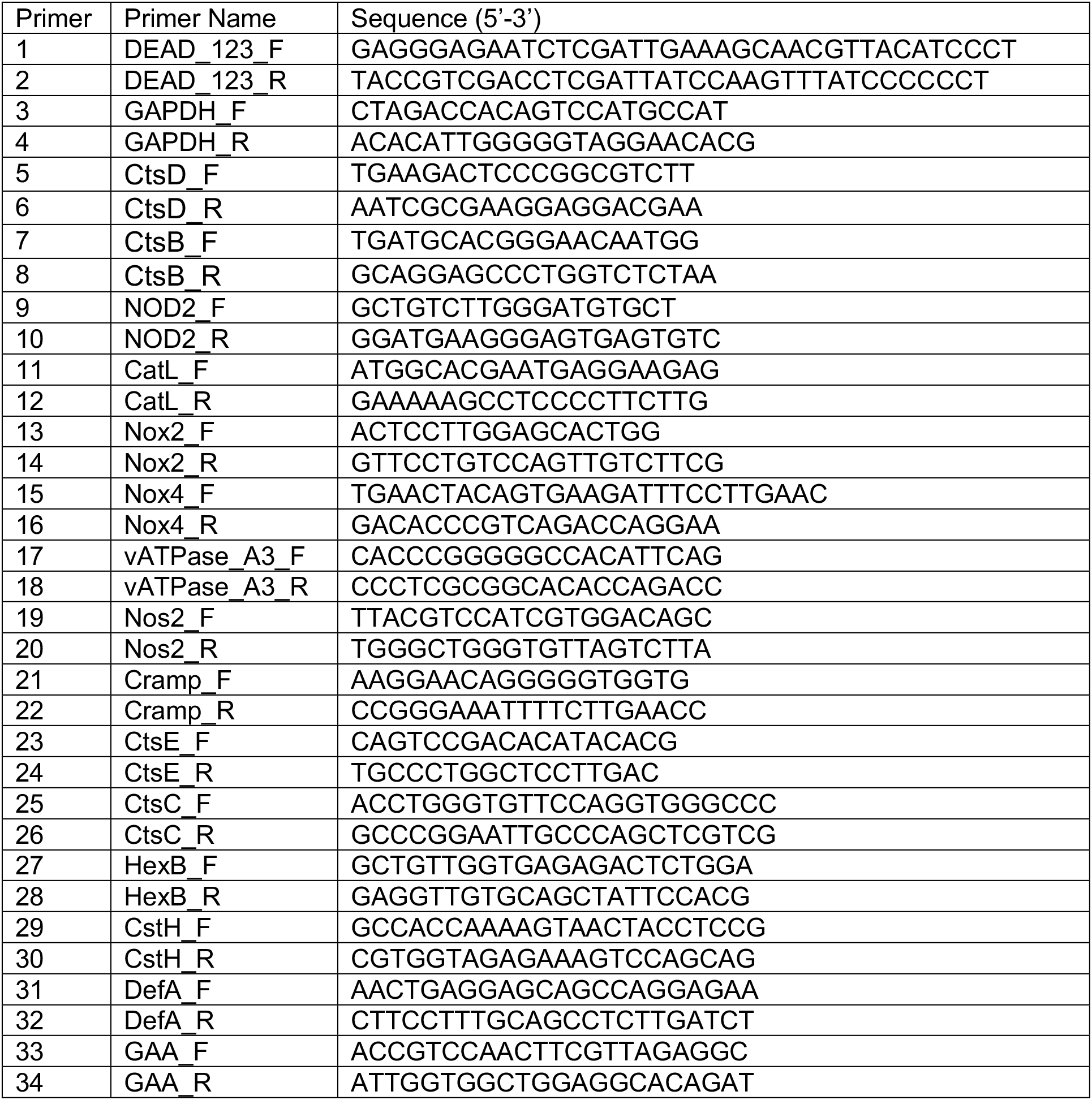
Primers used in this study.

## Author contributions

R.A.G.D.S., J.J.W., H.A., P.Y.C., K.K.Y., S.J., C.L., L.T.K.S, G.H. and M.V. performed experiments and analyzed data. R.A.G.D.S., H.A., C.L., J.C. and K.A.K. interpreted the results. K.A.K, J.C. and R.A.G.D.S.. designed the experiments and devised the project. R.A.G.D.S., J.C., and K.A.K. wrote the manuscript. All authors reviewed and approved the final manuscript.

## Data and materials availability

All data associated with this study are present in the paper or the Supplementary Materials.

## Acknowledgments

We thank Dr. Thomas Dean Watts and Dr. Claudia Jade Stocks for critical reading of the manuscript. We also thank Ms. Cheryl Neo Jia Yi and Mr. Muhaimin Bin Hasbullah for support with animal work.

## References

1. C. J. Murray, K. S. Ikuta, F. Sharara, L. Swetschinski, G. Robles Aguilar, A. Gray, C. Han, C. Bisignano, P. Rao, E. Wool, S. C. Johnson, A. J. Browne, M. G. Chipeta, F. Fell, S. Hackett, G. Haines-Woodhouse, B. H. Kashef Hamadani, E. A. P. Kumaran, B. McManigal, R. Agarwal, S. Akech, S. Albertson, J. Amuasi, J. Andrews, A. Aravkin, E. Ashley, F. Bailey, S. Baker, B. Basnyat, A. Bekker, R. Bender, A. Bethou, J. Bielicki, S. Boonkasidecha, J. Bukosia, C. Carvalheiro, C. Castañeda-Orjuela, V. Chansamouth, S. Chaurasia, S. Chiurchiù, F. Chowdhury, A. J. Cook, B. Cooper, T. R. Cressey, E. Criollo-Mora, M. Cunningham, S. Darboe, N. P. J. Day, M. de Luca, K. Dokova, A. Dramowski, S. J. Dunachie, T. Eckmanns, D. Eibach, A. Emami, N. Feasey, N. Fisher-Pearson, K. Forrest, D. Garrett, P. Gastmeier, A. Z. Giref, R. C. Greer, V. Gupta, S. Haller, A. Haselbeck, S. I. Hay, M. Holm, S. Hopkins, K. C. Iregbu, J. Jacobs, D. Jarovsky, F. Javanmardi, M. Khorana, N. Kissoon, E. Kobeissi, T. Kostyanev, F. Krapp, R. Krumkamp, A. Kumar, H. H. Kyu, C. Lim, D. Limmathurotsakul, M. J. Loftus, M. Lunn, J. Ma, N. Mturi, T. Munera-Huertas, P. Musicha, M. M. Mussi-Pinhata, T. Nakamura, R. Nanavati, S. Nangia, P. Newton, C. Ngoun, A. Novotney, D. Nwakanma, C. W. Obiero, A. Olivas-Martinez, P. Olliaro, E. Ooko, E. Ortiz-Brizuela, A. Y. Peleg, C. Perrone, N. Plakkal, A. Ponce-de-Leon, M. Raad, T. Ramdin, A. Riddell, T. Roberts, J. V. Robotham, A. Roca, K. E. Rudd, N. Russell, J. Schnall, J. A. G. Scott, M. Shivamallappa, J. Sifuentes-Osornio, N. Steenkeste, A. J. Stewardson, T. Stoeva, N. Tasak, A. Thaiprakong, G. Thwaites, C. Turner, P. Turner, H. R. van Doorn, S. Velaphi, A. Vongpradith, H. Vu, T. Walsh, S. Waner, T. Wangrangsimakul, T. Wozniak, P. Zheng, B. Sartorius, A. D. Lopez, A. Stergachis, C. Moore, C. Dolecek, M. Naghavi, Global burden of bacterial antimicrobial resistance in 2019: a systematic analysis. The Lancet 399, 629–655 (2022).

2. New report calls for urgent action to avert antimicrobial resistance crisis (available at https://www.who.int/news/item/29-04-2019-new-report-calls-for-urgent-action-to-avert-antimicrobial-resistance-crisis).

3. C. Y. Chiang, I. Uzoma, R. T. Moore, M. Gilbert, A. J. Duplantier, R. G. Panchal, Mitigating the impact of antibacterial drug resistance through host-directed therapies: Current progress, outlook, and challenges. mBio 9 (2018), doi:10.1128/MBIO.01932-17/ASSET/AF1059F5-51EA-4152-8BB3-FDBE7D97F121/ASSETS/GRAPHIC/MBO0011836790003.JPEG.

4. A. Giacometti, O. Cirioni, A. M. Schimizzi, M. S. del Prete, F. Barchiesi, M. M. D’Errico, E. Petrelli, G. Scalise, Epidemiology and microbiology of surgical wound infections. J Clin Microbiol 38, 918–922 (2000).

5. P. G. Bowler, B. I. Duerden, D. G. Armstrong, Wound microbiology and associated approaches to wound management. Clin Microbiol Rev 14, 244–269 (2001).

6. S. E. Dowd, Y. Sun, P. R. Secor, D. D. Rhoads, B. M. Wolcott, G. A. James, R. D. Wolcott, Survey of bacterial diversity in chronic wounds using Pyrosequencing, DGGE, and full ribosome shotgun sequencing. BMC Microbiology 8, 1–15 (2008).

7. L. J. Bessa, P. Fazii, M. di Giulio, L. Cellini, Bacterial isolates from infected wounds and their antibiotic susceptibility pattern: some remarks about wound infection. International Wound Journal 12, 47–52 (2015).

8. M. O. Ahmed, K. E. Baptiste, Vancomycin-Resistant Enterococci: A Review of Antimicrobial Resistance Mechanisms and Perspectives of Human and Animal Health. Microbial Drug Resistance 24, 590–606 (2018).

9. 2019 Antibiotic Resistance Threats Report | CDC (available at https://www.cdc.gov/drugresistance/biggest-threats.html#van).

10. H. S. Gold, Vancomycin-resistant enterococci: Mechanisms and clinical observations. Clinical Infectious Diseases 33, 210–219 (2001).

11. P. J. Stogios, A. Savchenko, Molecular mechanisms of vancomycin resistance. Protein Science 29, 654–669 (2020).

12. M. L. Faron, N. A. Ledeboer, B. W. Buchan, Resistance mechanisms, epidemiology, and approaches to screening for vancomycin-resistant Enterococcus in the health care setting. Journal of Clinical Microbiology 54, 2436–2447 (2016).

13. B. Y. Q. Tien, H. M. S. Goh, K. K. L. Chong, S. Bhaduri-Tagore, S. Holec, R. Dress, F. Ginhoux, M. A. Ingersoll, R. B. H. Williams, K. A. Kline, Enterococcus faecalis Promotes Innate Immune Suppression and Polymicrobial Catheter-Associated Urinary Tract Infection. Infect Immun 85 (2017), doi:10.1128/IAI.00378-17.

14. R. A. G. Da, S. Id, W. Hong Tay Id, F. Kiong, H. Id, F. Reinhart Tanoto, K. K. L. Chongid, P. Y. Choo, A. Ludwigid, K. A. Klineid, A. P. Hakansson, Ed. Enterococcus faecalis alters endo-lysosomal trafficking to replicate and persist within mammalian cells. PLOS Pathogens 18, e1010434 (2022).

15. K. K. L. Chong, W. H. Tay, B. Janela, A. M. H. Yong, T. H. Liew, L. Madden, D. Keogh, T. M. S. Barkham, F. Ginhoux, D. L. Becker, K. A. Kline, Enterococcus faecalis Modulates Immune Activation and Slows Healing During Wound Infection. The Journal of Infectious Diseases 216, 1644 (2017).

16. F. C. Fang, Antimicrobial reactive oxygen and nitrogen species: concepts and controversies. Nat Rev Microbiol 2, 820–832 (2004).

17. J. M. Slauch, How does the oxidative burst of macrophages kill bacteria? Still an open question. Mol Microbiol 80, 580 (2011).

18. J. R. Sheldon, E. P. Skaar, Metals as phagocyte antimicrobial effectors. Curr Opin Immunol 60, 1 (2019).

19. G. L. Lukacs$$ll, O. D. Rotstein, & S. Grinstein+, Phagosomal acidification is mediated by a vacuolar-type H(+)-ATPase in murine macrophages. THE JOURNAL. OF BIOLOGICAL CHEMISTRY 265, 21099–21107 (1990).

20. E. Uribe-Querol, C. Rosales, Phagocytosis: Our Current Understanding of a Universal Biological Process. Frontiers in Immunology 11, 1066 (2020).

21. M. Orecchioni, Y. Ghosheh, A. B. Pramod, K. Ley, Macrophage polarization: Different gene signatures in M1(Lps+) vs. Classically and M2(LPS-) vs. Alternatively activated macrophages. Frontiers in Immunology 10, 1084 (2019).

22. P. J. Murray, Macrophage Polarization. http://dx.doi.org/10.1146/annurev-physiol-022516-034339 79, 541–566 (2017).

23. C. D. Mills, K. Kincaid, J. M. Alt, M. J. Heilman, A. M. Hill, M-1/M-2 macrophages and the Th1/Th2 paradigm. J Immunol 164, 6166–6173 (2000).

24. U. Theuretzbacher, K. Outterson, A. Engel, A. Karlén, The global preclinical antibacterial pipelineNature Reviews Microbiology 18, 275–285 (2020).

25. G. Hu, Y. Su, B. H. Kang, Z. Fan, T. Dong, D. R. Brown, J. Cheah, K. D. Wittrup, J. Chen, High-throughput phenotypic screen and transcriptional analysis identify new compounds and targets for macrophage reprogramming. Nat Commun 12 (2021), doi:10.1038/S41467-021-21066-X.

26. G. Hu, Y. Su, B. H. Kang, Z. Fan, T. Dong, D. R. Brown, J. Cheah, K. D. Wittrup, J. Chen, High-throughput phenotypic screen and transcriptional analysis identify new compounds and targets for macrophage reprogramming. Nature Communications 12 (2021), doi:10.1038/s41467-021-21066-x.

27. M. J. Rybak, B. M. Lomaestro, J. C. Rotschafer, R. C. Moellering, W. A. Craig, M. Billeter, J. R. Dalovisio, D. P. Levine, Vancomycin Therapeutic Guidelines: A Summary of Consensus Recommendations from the Infectious Diseases Society of America, the American Society of Health-System Pharmacists, and the Society of Infectious Diseases Pharmacists. Clinical Infectious Diseases 49, 325–327 (2009).

28. R. F. Novak, E. D. Kharasch, Mitoxantrone: propensity for free radical formation and lipid peroxidation--implications for cardiotoxicity. Invest New Drugs 3, 95–99 (1985).

29. F. Baquero, B. R. Levin, Proximate and ultimate causes of the bactericidal action of antibiotics. Nature Reviews Microbiology 2020 19:2 19, 123–132 (2020).

30. C. Watanakunakorn, Mode of action and in-vitro activity of vancomycin. J Antimicrob Chemother 14 Suppl D, 7–18 (1984).

31. D. H. Bell, Characterization of the fluorescence of the antitumor agent, mitoxantrone. Biochim Biophys Acta 949, 132–137 (1988).

32. P. Redder, S. Hausmann, V. Khemici, H. Yasrebi, P. Linder, Bacterial versatility requires DEAD-box RNA helicases. FEMS Microbiology Reviews 39, 392–412 (2015).

33. M. Benoit, B. Desnues, J.-L. Mege, Macrophage Polarization in Bacterial Infections. The Journal of Immunology 181, 3733–3739 (2008).

34. Q. Huang, J. Hou, P. Yang, J. Yan, X. Yu, Y. Zhuo, S. He, F. Xu, Antiviral activity of mitoxantrone dihydrochloride against human herpes simplex virus mediated by suppression of the viral immediate early genes. BMC Microbiology 19, 1–9 (2019).

35. P. F. Chan, V. Srikannathasan, J. Huang, H. Cui, A. P. Fosberry, M. Gu, M. M. Hann, M. Hibbs, P. Homes, K. Ingraham, J. Pizzollo, C. Shen, A. J. Shillings, C. E. Spitzfaden, R. Tanner, A. J. Theobald, R. A. Stavenger, B. D. Bax, M. N. Gwynn, Structural basis of DNA gyrase inhibition by antibacterial QPT-1, anticancer drug etoposide and moxifloxacin. Nature Communications 2015 6:1 6, 1–13 (2015).

36. E. J. Fox, Mechanism of action of mitoxantrone. Neurology 63, S15–S18 (2004).

37. R. J. Worthington, C. Melander, Combination Approaches to Combat Multi-Drug Resistant Bacteria. Trends Biotechnol 31, 177 (2013).

38. M. L. Faron, N. A. Ledeboer, B. W. Buchan, Resistance mechanisms, epidemiology, and approaches to screening for vancomycin-resistant Enterococcus in the health care setting. Journal of Clinical Microbiology 54, 2436–2447 (2016).

39. W. H. Tay, R. A. G. da Silva, F. K. Ho, K. K. L. Chong, A. Ludwig, K. A. Kline, Enterococcus faecalis persists and replicates within epithelial cells in vitro and in vivo during wound infection. bioRxiv, 2021.09.16.460717 (2021).

40. K. J. I. Thorne, R. C. Oliver, A. J. Barrett, Lysis and killing of bacteria by lysosomal proteinases. Infection and Immunity 14, 555 (1976).

41. A. Reis-mendes, J. L. Dores-sousa, A. I. Padrão, M. Duarte-araújo, J. A. Duarte, V. Seabra, S. Gonçalves-monteiro, F. Remião, F. Carvalho, E. Sousa, M. L. Bastos, V. M. Costa, Inflammation as a Possible Trigger for Mitoxantrone-Induced Cardiotoxicity: An In Vivo Study in Adult and Infant Mice. Pharmaceuticals (Basel) 14 (2021), doi:10.3390/PH14060510.

42. L. Fischer-Riepe, N. Daber, J. Schulte-Schrepping, B. C. Véras De Carvalho, A. Russo, M. Pohlen, J. Fischer, A. I. Chasan, M. Wolf, T. Ulas, S. Glander, C. Schulz, B. Skryabin, A. Wollbrink, Dipl-Ing, N. Steingraeber, C. Stremmel, M. Koehle, F. Gärtner, S. Vettorazzi, D. Holzinger, J. Gross, F. Rosenbauer, M. Stoll, S. Niemann, J. Tuckermann, J. L. Schultze, J. Roth, K. Barczyk-Kahlert, CD163 expression defines specific, IRF8-dependent, immune-modulatory macrophages in the bone marrow. Journal of Allergy and Clinical Immunology 146, 1137–1151 (2020).

43. D. Parker, CD80/CD86 signaling contributes to the proinflammatory response of Staphylococcus aureus in the airway. Cytokine 107, 130 (2018).

44. M. C. Marchitto, C. A. Dillen, H. Liu, R. J. Miller, N. K. Archer, R. v. Ortines, M. P. Alphonse, A. I. Marusina, A. A. Merleev, Y. Wang, B. L. Pinsker, A. S. Byrd, I. D. Brown, A. Ravipati, E. Zhang, S. S. Cai, N. Limjunyawong, X. Dong, M. R. Yeaman, S. I. Simon, W. Shen, S. K. Durum, R. L. O’Brien, E. Maverakis, L. S. Miller, Clonal Vγ6+Vδ4+ T cells promote IL-17–mediated immunity against Staphylococcus aureus skin infection. Proc Natl Acad Sci U S A 166, 10917–10926 (2019).

45. C. A. Dillen, B. L. Pinsker, A. I. Marusina, A. A. Merleev, O. N. Farber, H. Liu, N.K. Archer, D. B. Lee, Y. Wang, R. v. Ortines, S. K. Lee, M. C. Marchitto, S. S. Cai, A. G. Ashbaugh, L. S. May, S. M. Holland, A. F. Freeman, L. G. Miller, M. R. Yeaman, S. I. Simon, J. D. Milner, E. Maverakis, L. S. Miller, Clonally expanded γδ T cells protect against Staphylococcus aureus skin reinfection. The Journal of Clinical Investigation 128, 1026 (2018).

46. L. J. Juttukonda, W. N. Beavers, D. Unsihuay, K. Kim, G. Pishchany, K. J. Horning, A. Weiss, H. Al-Tameemi, J. M. Boyd, G. A. Sulikowski, A. B. Bowman, E. P. Skaara, A small-molecule modulator of metal homeostasis in gram-positive pathogens. mBio 11, 1–22 (2020).

47. L. Cui, Y. H. Lee, T. L. Thein, J. Fang, J. Pang, E. E. Ooi, Y. S. Leo, C. N. Ong, S. R. Tannenbaum, Serum Metabolomics Reveals Serotonin as a Predictor of Severe Dengue in the Early Phase of Dengue Fever. PLoS Negl Trop Dis 10 (2016), doi:10.1371/JOURNAL.PNTD.0004607.

48. B. Luna, V. Trebosc, B. Lee, M. Bakowski, A. Ulhaq, J. Yan, P. Lu, J. Cheng, T. Nielsen, J. Lim, W. Ketphan, H. Eoh, C. McNamara, N. Skandalis, R. She, C. Kemmer, S. Lociuro, G. E. Dale, B. Spellberg, A nutrient-limited screen unmasks rifabutin hyperactivity for extensively drug-resistant Acinetobacter baumannii. Nature Microbiology 5, 1134–1143 (2020).

49. K. L. Palmer, A. Daniel, C. Hardy, J. Silverman, M. S. Gilmore, Genetic basis for daptomycin resistance in enterococci. Antimicrobial Agents and Chemotherapy 55, 3345–3356 (2011).

50. H. v. Nielsen, P. S. Guiton, K. A. Kline, G. C. Port, J. S. Pinkner, F. Neiers, S. Normark, B. Henriques-Normark, M. G. Caparon, S. J. Hultgren, The metal ion-dependent adhesion site motif of the Enterococcus faecalis EbpA pilin mediates pilus function in catheter-associated urinary tract infection. mBio 3 (2012), doi:10.1128/MBIO.00177-12.

51. S. Manzanero, Generation of mouse bone marrow-derived macrophages. Methods Mol Biol 844, 177–181 (2012).

52. K. J. Livak, T. D. Schmittgen, Analysis of relative gene expression data using real-time quantitative PCR and the 2(-Delta Delta C(T)) Method. Methods 25, 402–408 (2001).

53. K. Eliceiri, C. A. Schneider, W. S. Rasband, K. W. Eliceiri, NIH Image to ImageJ: 25 years of image analysis HISTORICAL commentary NIH Image to ImageJ: 25 years of image analysis. Nature Methods 9, 671–675 (2012).

54. G. M. Dunny, B. L. Brown, D. B. Clewell, Induced cell aggregation and mating in Streptococcus faecalis: evidence for a bacterial sex pheromone. Proc Natl Acad Sci U S A 75, 3479–3483 (1978).

55. A. Bourgogne, D. A. Garsin, X. Qin, K. v. Singh, J. Sillanpaa, S. Yerrapragada, Y. Ding, S. Dugan-Rocha, C. Buhay, H. Shen, G. Chen, G. Williams, D. Muzny, A. Maadani, K. A. Fox, J. Gioia, L. Chen, Y. Shang, C. A. Arias, S. R. Nallapareddy, M. Zhao, V. P. Prakash, S. Chowdhury, H. Jiang, R. A. Gibbs, B. E. Murray, S. K. Highlander, G. M. Weinstock, Large scale variation in Enterococcus faecalis illustrated by the genome analysis of strain OG1RF. Genome Biology 9, 1–16 (2008).

56. M. Totsika, S. A. Beatson, S. Sarkar, M. D. Phan, N. K. Petty, N. Bachmann, M. Szubert, H. E. Sidjabat, D. L. Paterson, M. Upton, M. A. Schembri, Insights into a multidrug resistant Escherichia coli pathogen of the globally disseminated ST131 lineage: genome analysis and virulence mechanisms. PLoS One 6 (2011), doi:10.1371/JOURNAL.PONE.0026578.

57. S. L. Chen, C. S. Hung, J. Xu, C. S. Reigstad, V. Magrini, A. Sabo, D. Blasiar, T. Bieri, R. R. Meyer, P. Ozersky, J. R. Armstrong, R. S. Fulton, J. P. Latreille, J. Spieth, T. M. Hooton, E. R. Mardis, S. J. Hultgren, J. I. Gordon, Identification of genes subject to positive selection in uropathogenic strains of Escherichia coli: a comparative genomics approach. Proc Natl Acad Sci U S A 103, 5977–5982 (2006).

58. L. K. McDougal, C. D. Steward, G. E. Killgore, J. M. Chaitram, S. K. McAllister, F. C. Tenover, Pulsed-Field Gel Electrophoresis Typing of Oxacillin-Resistant Staphylococcus aureus Isolates from the United States: Establishing a National Database. Journal of Clinical Microbiology 41, 5113 (2003).

59. J. C. O. T. I. C. O. S. O. P. None, The type species of the genus Salmonella Lignieres 1900 is Salmonella enterica (ex Kauffmann and Edwards 1952) Le Minor and Popoff 1987, with the type strain LT2T, and conservation of the epithet enterica in Salmonella enterica over all earlier epithets that may be applied to this species. Opinion 80. Int J Syst Evol Microbiol 55, 519–520 (2005).

60. M. Hentzer, K. Riedel, T. B. Rasmussen, A. Heydorn, J. B. Andersen, M. R. Parsek, S. A. Rice, L. Eberl, S. Molin, N. Høiby, S. Kjelleberg, M. Givskov, Inhibition of quorum sensing in Pseudomonas aeruginosa biofilm bacteria by a halogenated furanone compound. Microbiology (Reading) 148, 87–102 (2002).

61. X. Zhang, V. de Maat, A. M. Guzmán Prieto, T. K. Prajsnar, J. R. Bayjanov, M. de Been, M. R. C. Rogers, M. J. M. Bonten, S. Mesnage, R. J. L. Willems, W. van Schaik, RNA-seq and Tn-seq reveal fitness determinants of vancomycin-resistant Enterococcus faecium during growth in human serum. BMC Genomics 18, 1–12 (2017).

62. M. M. C. Lam, T. Seemann, D. M. Bulach, S. L. Gladman, H. Chen, V. Haring, R. J. Moore, S. Ballard, M. L. Grayson, P. D. R. Johnson, B. P. Howden, T. P. Stineara, Comparative Analysis of the First Complete Enterococcus faecium Genome. Journal of Bacteriology 194, 2334 (2012).

63. J. Top, S. Arredondo-Alonso, A. C. Schürch, S. Puranen, M. Pesonen, J. Pensar, R. J. L. Willems, J. Corander, Genomic rearrangements uncovered by genome-wide co-evolution analysis of a major nosocomial pathogen, Enterococcus faecium. Microbial Genomics 6, 1–8 (2020).

